# Decoding Clade-Specific bNAb–HIV-1 Env Interactions Using Machine Learning

**DOI:** 10.64898/2026.10.14.759022

**Authors:** Yogendra Kumar, Shiang-Tai Lin, Narendra M. Dixit, Prabal K. Maiti

## Abstract

Human immunodeficiency virus type 1 (HIV-1) exhibits extensive genetic diversity, posing a major challenge to AIDS vaccine development. Broadly neutralizing antibodies (bNAbs) isolated from HIV-1-infected individuals offer valuable templates for designing broadly protective immunogens. Here, structural and energetic descriptors extracted from experimentally resolved bNAb/HIV-1 Envelope (Env) protein-protein complexes were used to develop machine-learning (ML) models predicting binding free energies (ΔG) and dissociation constants (*K*_*D*_) across multiple HIV-1 clades. Among the algorithms evaluated, extreme gradient boosting (XGBoost) achieved the best predictive performance, with training and test *R*^2^ values of 0.92 and 0.658, respectively. Applying the optimized model to HIV-1 datasets revealed clade-dependent binding patterns, with clade B showing the greatest variability and clade G exhibiting signatures associated with reduced neutralization potency. Predicted ΔG of different bNAb classes followed the trend gp120-gp41 interface ≈ fusion domain *>* CD4 binding site (CD4bs) *>* V1V2 loop ≈ V3-glycan, consistent with experimentally reported affinity profiles of benchmark bNAbs. Clade C exhibited intermediate ΔG with PGT121, VRC01, and 10-1074 maintaining high-affinity interactions, whereas PG9 and 2G12 show reduced binding affinity. These findings offer quantitative insight into clade-specific bNAb binding energetics and demonstrate the potential of machine-learning approaches to guide HIV-1 immunogen design and vaccine development.

## Introduction

Human immunodeficiency virus type 1 (HIV-1) poses a serious threat to global health, as it is the causative agent of acquired immune deficiency syndrome (AIDS).^1^ Although substantial worldwide efforts have been made to control the spread of HIV-1, new infections and AIDS-related deaths remain a major public health concern.^2,3^ The World Health Organization estimates that 40.8 million people were living with HIV-1 globally at the end of 2024, and approximately 630,000 people died from AIDS-related causes that year.^4,5^ Sub-Saharan Africa continues to bear the greatest burden of the epidemic, accounting for more than half of all people living with HIV worldwide. Within this region, women and girls are disproportionately affected, making up 63% of new HIV infections.^5^

The HIV-1 Envelope (Env) glycoprotein, gp160, forms a trimeric structure made up of three gp120 surface subunits and three gp41 transmembrane subunits. Together, they create a metastable complex that is essential for viral entry.^6–8^ This gp120-gp41 assembly enables viral entry by binding to host cell receptors and driving membrane fusion.^9–16^ Once binding occurs, gp120 undergoes a series of conformational changes that trigger gp41 to initiate the fusion process.^17,18^ After fusion with the host cell is complete, the virus enters the post-fusion stage, where viral replication begins.^18^

Antiretroviral therapy (ART) remains the cornerstone of HIV-1 management, substantially reducing morbidity and mortality by suppressing viral replication and preserving immune function.^3,19,20^ However, individuals on prolonged ART may develop drug resistance over time.^19,20^ Broadly neutralizing antibodies (bNAbs) have emerged as a highly effective approach to HIV-1 treatment.^21–24^

During chronic HIV-1 infection, a subset of individuals develops bNAbs that can effectively neutralize diverse viral strains across multiple clades. These antibodies, isolated from the sera of infected individuals target conserved regions of the HIV-1 Env glycoprotein.^25^ bNAbs neutralize the virus by binding specific epitopes on Env, thereby blocking receptor engagement or the fusion process. Six major epitope regions have been identified as bNAb targets: the CD4 binding site (CD4bs), the V1/V2-glycan apex, the V3-glycan region, the gp120/gp41 interface, the silent face of gp120, and the membrane-proximal external region (MPER) of gp41.^26^ These epitopes, mapped for HIV-1 subtype/clade C, are shown in Figure 1. Recent studies show that combining bNAbs targeting different epitopes enhances neutralization breadth and potency, reduces the risk of viral escape, and supports sustained long-term suppression.^27^

**Figure 1:**
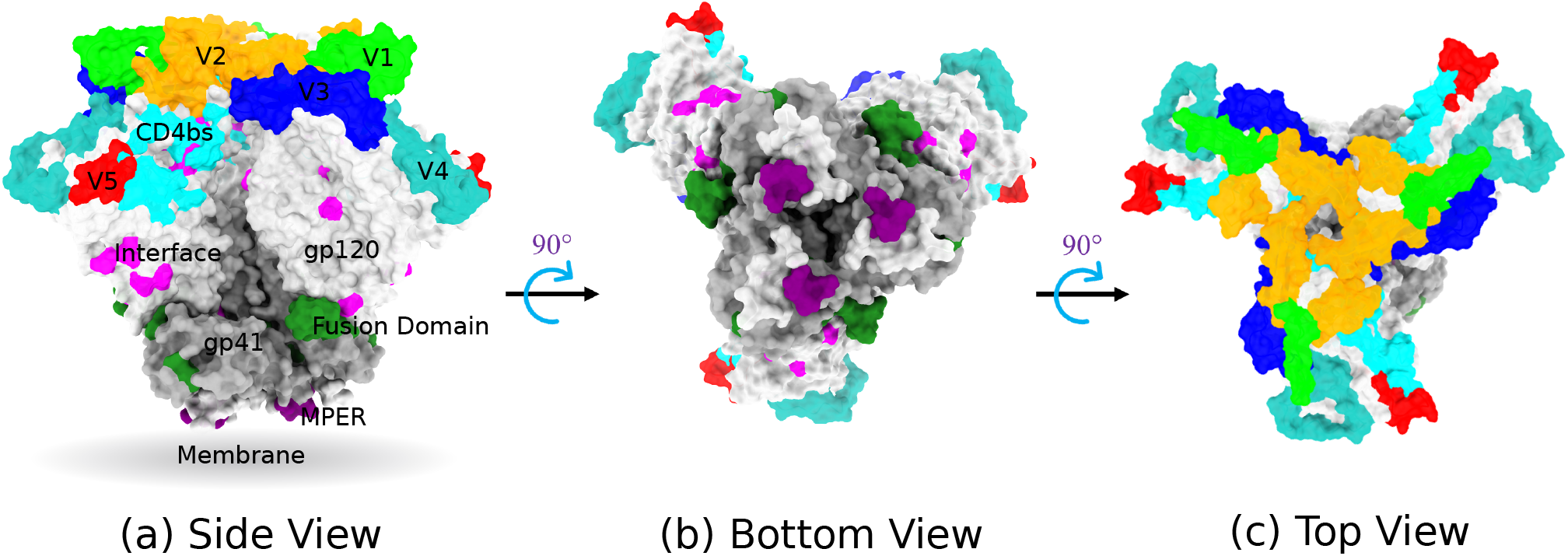
Snapshot of the HIV-1 Env trimer subtype clade C (PDB ID: 8SAU) in the different colors of the major bNAbs targeting epitopes in (a) side view, (b) bottom view, and (c) top view. Color Code: CD4bs in cyan, V1 loops in lime, V2 loops in orange, V3 loops in blue, V4 loops in light sea green, V5 loops in red, gp41/gp120 interface in magenta, fusion domains in green, and MPER region in purple. Env gp120 and gp41 are shown in light and dark grey, respectively. Residue numbers for all the epitopes are based on the HxB2 sequence.

The gp120 subunit of the HIV-1 Env glycoprotein is among the most heavily glycosylated proteins known, with N-linked glycans accounting for roughly half of its molecular mass.^28–31^ These glycans form a dense, heterogeneous shield that masks underlying protein epitopes and limits antibody access.^30,31^ Because this glycan shield is dynamic, it can evolve under immune pressure, allowing the virus to escape host antibody responses. Nevertheless, some bNAbs have evolved to recognize glycan-dependent epitopes, such as those in the V1/V2 and V3 regions, by simultaneously engaging glycan moieties and adjacent protein surfaces.^21,32,33^

Glycans play a pivotal role in modulating the binding affinity and specificity of bNAbs to their respective epitopes on the HIV-1 Env glycoprotein. Many bNAbs rely on specific glycan moieties for high-affinity interactions and effective neutralization.^34^ The structural diversity and spatial arrangement of these glycans contribute significantly to the antigenic landscape of Env, influencing both antibody accessibility and neutralization breadth.^34,35^

Recently, Basu et al. used microscale thermophoresis to measure the binding affinity (*K*_*d*_) of bNAbs and monoclonal antibodies targeting different epitopes on the Env glycoprotein.^36^ Among these, CD4bs-specific bNAbs such as VRC01 and 3BNC117 showed the tightest binding to trimeric HIV-1 Env proteins,^36,37^ whereas bNAbs targeting the V1/V2, V3, and MPER epitopes showed considerably weaker binding, with *K*_*d*_ values ranging from less than 1 nM to 40 nM.^36–38^ In a related study, Alam et al. found that deleting 11 N-terminal amino acids enhanced the immunogenicity and antigenicity of the AIDSVAX HIV-1 vaccine for subtype B/E. For the V1/V2 epitope targeted by the bNAb PG9, they reported *K*_*d*_ values of 183 nM, 48.1 nM, and 53.6 nM for gp120 from AV144, gDΔ11 gp120 from A244, and E.A244Δ11 gp120, respectively.^39^ Notably, the AIDSVAX vaccine trial in Thailand demonstrated 31.2% efficacy in preventing HIV-1 acquisition among heterosexuals,^40^ prompting broader discussion of whether Env-based vaccines can reliably induce protective, infection-preventing antibodies.^38^

Building on this, Bontempo et al. reported that the bNAbs PG9, PGT145, VRC01, and PGT121 bind much more strongly to the HIV-1 Env glycoprotein of subtype BG505 SOSIP.664, with *K*_*d*_ values of 2.1 nM, 2.57 nM, ≤ 2.1 × 10^*−*3^ nM, and 0.15 nM, respectively.^37^ Extending this work, Garces et al. focused on the high-mannose patch region of recombinant BG505 SOSIP.664 Env and characterized the binding affinity of the PGT121 family of bNAbs.^41^

Beyond antibody studies, molecular dynamics (MD) simulations have also been used to explore therapeutic strategies: several studies have shown the potential of dendrimers to target the CD4bs.^42–44^ Separately, Feng et al. examined the interaction between the VRC34.01 antibody and the HIV-1 Env fusion peptide, finding that mutations at key binding residues—such as Ile515-significantly reduce antibody affinity, highlighting how sensitive epitope recognition is to sequence variation.^45^ Similarly, PGT121 and 10-1074, both capable of recognizing glycan-shielded epitopes within the V3 loop of Env,^32,46^ target the N332 glycan and its surrounding region; however, structural and biophysical analyses show that 10-1074 binds with higher affinity than PGT121, as reflected in its lower *K*_*d*_.^32^

In a complementary line of work, Garg et al. developed a model explaining how passive immunization (PI) with high-affinity antibodies can enhance endogenous antibody production, combining experimental data with strategies to optimize PI protocols for stronger immune responses.^47^ Meanwhile, other studies highlight a key clinical challenge: HIV-1 variants resistant to VRC01 can persist within the latent reservoir-a pool of dormant infected cells unaffected by ART. When ART is discontinued and VRC01 is administered, these resistant strains can reactivate and replicate, leading to treatment failure.^48,49^ At the structural level, gp41 behaves differently before and after fusion: in the pre-fusion state, it interacts with lipids to form a stalk, while simple membranes resist fusion;^50^ post-fusion, gp41 binds strongly to the inhibitor C34 but only weakly to T20.^51^

Although binding affinity (*K*_*d*_) is a key determinant of the interaction strength between bNAbs and the HIV-1 Env glycoprotein, it remains relatively underexplored compared to traditional neutralization metrics such as IC50. To address this gap, we employed machine-learning regression models to predict *K*_*d*_ values of bNAbs across diverse HIV-1 clades, offering quantitative insights into clade-dependent anti-body–Env interactions.

## Materials and methods

### Data Collection and Curation

We obtained 3D structures of clinical trial bNAbs (Table S1) and HIV-1 Env complexes from the Protein Data Bank (www.RCSB.org).^52^ HIV-1 Env trimer structures for clades A (5FYL), B (5FYK), C (8SAU), and G (5FYJ) were taken from our previous study.^24^ Each bNAb/HIV-1 complex was superimposed onto the corresponding subtype structure, after which the original PDB-derived HIV-1 unit was removed to generate the bNAb bound to the HIV-1 Env trimer (schematic in Figure S1, SI). The resulting structures were curated into uniform chain labels for HIV-1 (A, B, C, G) and bNAbs (H, L) across all subtype complexes. IC50 values for the corresponding bNAbs were collected from the literature.^36,53,54^

For model training and validation, we used benchmark protein-protein complex data from the PDBbind+ database,^55^ which contains 2,852 entries, each with a 3D structure (PDB format) and an experimentally determined binding affinity value (*K*_*d*_). The distribution of the target variable after curation is shown in Figure S2 (SI).

### MD Simulations

All-atom molecular dynamics simulations were performed using GROMACS v2022.3^56^ for bNAb complexes with non-glycosylated trimeric HIV-1 Env from various clades: 5FYL (clade A), 5FYK (clade B), 8SAU (clade C), and 5FYJ (clade G). Topology and coordinate files for each non-glycosylated Env-bNAb complex were generated using the CHARMM36 force field.^57^ Each complex was placed in an octahedral box with a minimum distance of 0.8 nm from the box edges and solvated using the TIP3P water model.^58^ All systems were added 150 mM NaCl mimic physiological ionic strength.

Energy minimization was carried out in two stages using the steepest-descent method: first, 10,000 steps with the trimer’s heavy atoms restrained (force constant of 500 kJ mol^*−*1^*nm*^*−*2^) to remove steric clashes, followed by an additional 10,000 steps without restraints. This was followed by NVT equilibration for 500 ps, during which the trimer’s heavy atoms remained restrained at the same force constant. Temperature was maintained at 300 K using the Nose-Hoover thermostat^59,60^ with a coupling constant of 1.0 ps, while isotropic pressure was controlled using the Parrinello–Rahman barostat^61^ with a coupling constant of 5 ps. The LINCS algorithm was used to constrain hydrogen bonds,^62^ non-bonded interactions were computed using the Verlet cutoff scheme (r = 1.2 nm), and long-range electrostatics were treated with the Particle Mesh Ewald (PME) method.^63,64^

Finally, a 20 ns unrestrained NPT production simulation was performed for each trimer, with trajectory frames saved every 5,000 steps. From the last 1 ns of each trajectory, 10 frames were extracted, and structural descriptors were computed from each frame to determine accurate binding affinities for the protein-protein complexes.

### Descriptors for Protein-Protein Complexes

The PDBbind+ v.2020R1 dataset^65^ contains 2,852 protein-protein complexes, many with multiple chains under a single PDB ID. We retained only the chains relevant to binding affinity, removing the rest using a ChimeraX script,^66^ yielding 2,485 curated structures for descriptor calculation.

Structural descriptors were selected to capture the contributions of charged, polar, apolar, hydrophilic, and hydrophobic amino acids at the protein-protein interface. For each of the 2,485 complexes, we calculated buried surface area (BSA), interchain contacts (ICs), and percent non-interacting surface (NIS) for charged-charged (CC), polar-polar (PP), and apolar-apolar (AP) residue pairs. All structural descriptors were computed using in-house Python code built on MDAnalysis.^67^ Interface hydrogen bonds were computed using HBPLUS,^68^ while energetic features were derived using HADDOCK,^69,70^ FoldX,^71^ and Rosetta.^72^ Correlations among all extracted features are shown in Figure 4.

The following equations were used to calculate the total BSA, NIS, and %NIS:

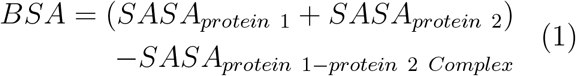

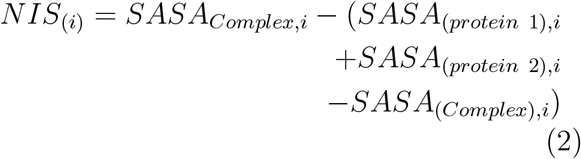

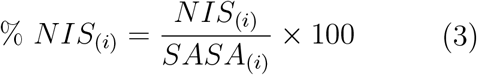

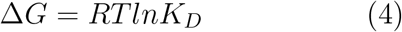

The descriptors BSA CC, BSA AA, and BSA PP were calculated using Eq. 1, corresponding to charged-charged, apolar-apolar, and polar-polar amino acid interactions within the protein-protein complexes, respectively. Similarly, %NIS CC, %NIS AA, and %NIS PP were calculated using Eqs. 2 and 3, where *i* denotes the charged-charged (CC), apolar-apolar (AA), or polar-polar (PP) interaction class. Binding free energy (ΔG, kcal mol^*−*1^) was calculated using Eq. 4, where *R* is the gas constant (0.001987 kcal mol^*−*1^K^*−*1^), *T* is the temperature (300 K), and *K*_*D*_ is the dissociation constant in *nM*. All features used in this study are summarized in Table S2 (Supporting Information).

### Model Building

To capture the complex relationships within the dataset and ensure robust predictive performance, four machine learning architectures were evaluated: Extreme Gradient Boosting (XGBoost),^73^ Random Forest Regression (RF),^74^ Support Vector Regression (SVR),^75^ and Decision Tree Regression (DT).^76^ A single Decision Tree served as the baseline model for its interpretability and non-parametric nature. To reduce the high variance typical of individual trees, a Random Forest ensemble was used, leveraging bootstrap aggregation to improve generalization. Support Vector Regression was chosen for its effectiveness in high-dimensional spaces and its ability to define a flexible decision boundary via the epsilon-insensitive loss function. Finally, XGBoost-a scalable, end-to-end gradient boosting framework-was used to capture complex non-linear patterns through its regularization and iterative optimization process.

Together, this diverse set of algorithms enables a comprehensive comparison across traditional tree-based methods, kernel-based optimization, and state-of-the-art boosting frameworks. The XGBoost model was trained with optimized hyperparameters: a maximum tree depth of 6, a learning rate of 0.05, 1000 estimators, subsampling and column-sampling ratios of 0.7, and an L2 regularization parameter (*λ*) of 10.0, balancing model complexity and generalization performance.

### Model Evaluation

The predictive performance and generalization ability of the four regression models (DT, RF, SVR, and XGBoost) were evaluated using three primary metrics: mean absolute error (MAE), root mean squared error (RMSE), and the coefficient of determination (*R*^2^). The equations used for these evaluation metrics are given below:

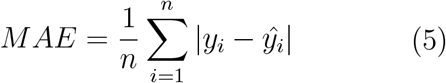

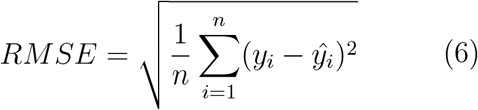

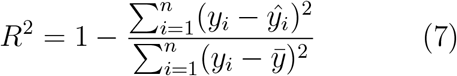

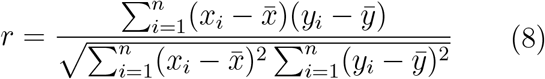

where *n, y*_*i*_, and *ŷ*_*i*_ are the number of observations, true value, and predicted value, respectively. 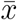 is the mean of the true values, and 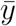 is the mean of the predicted values. *MAE* (Eq. 5) provides an intuitive measure of average error magnitude, while *RMSE* (Eq. 6) penalizes larger deviations more heavily, offering insight into model variance. *R*^2^ (Eq. 7) quantifies the proportion of variance in the dependent variable explained by the predictors, serving as a benchmark for overall goodness of fit. Additionally, the Pearson correlation coefficient (*r*) (Eq. 8) was used to assess model performance on the test set.

### Cross Validation

To ensure generalizability and minimize overfitting, the complete dataset of 2,445 entries was first split into a model-development set and an independent test set using a 75:25 ratio, yielding 1,833 and 612 entries, respectively. Within the model-development set, 184 entries were set aside for internal validation, while the remaining 1,649 entries were used for training and cross-validation.

A 5-fold cross-validation strategy was applied exclusively to the training subset. The 1,649 training entries were divided into five roughly equal folds; in each iteration, four folds were used for model fitting and the remaining fold for validation, with this process repeated five times so that each fold served as the validation set once. The average *R*^2^, *RMSE*, and *MAE* across the five iterations guided model selection and hyperparameter optimization.

The independent test set played no role in training, validation, or cross-validation and was reserved solely for final performance evaluation. This design enabled robust model optimization while preserving an unbiased assessment of generalization performance. The complete machine-learning workflow is summarized in Figure 2.

**Figure 2:**
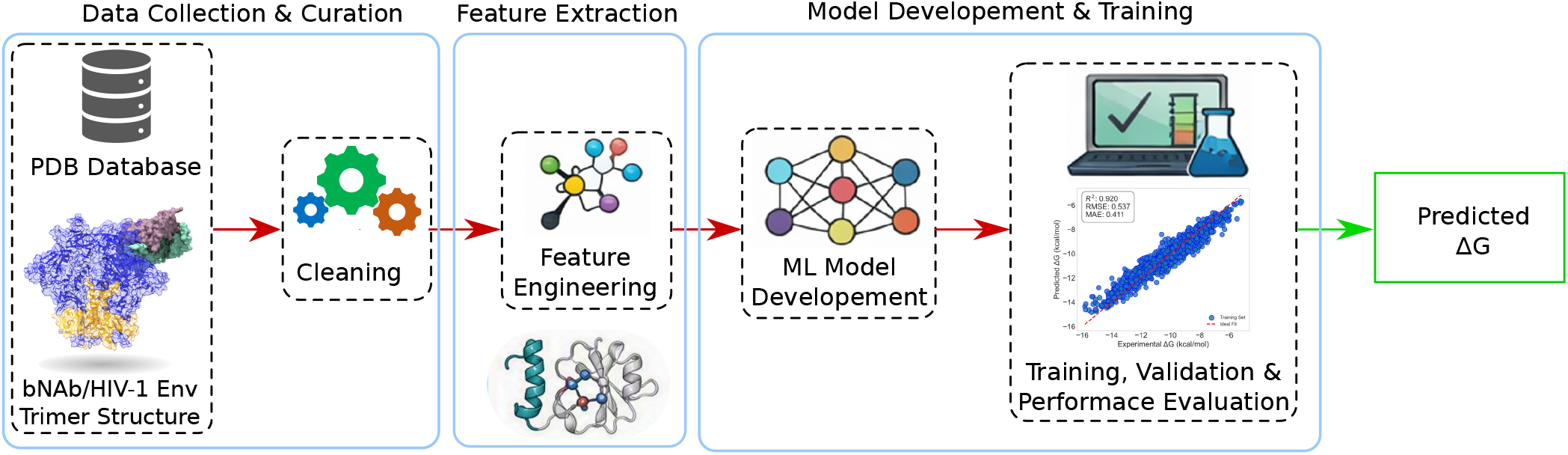
Machine Learning workflow used to collect, curate data and predict the binding affinity of bNAb/HIV-1 Env trimer complex.

## Results and Discussion

### Model Performance

Four supervised regression algorithms were evaluated: XGBoost, RF, SVR, and DT. Model performance was assessed using the coefficient of determination (*R*^2^), root mean square error (*RMSE*), and mean absolute error (*MAE*) as evaluation metrics (Table 1). Figure 3 presents comparison of the XGBoost-predicted binding energies (ΔG) with experimentally determined values for the training set (Figure 3(a)) and the combined test and validation sets (Figure 3(b)). In both panels, the dashed red diagonal line represents the locus of the ideal prediction (y = x), and the inset statistics quantify model performance through the coefficient of determination (*R*^2^), root mean square error (*RMSE*, kcal mol^*−*1^), and mean absolute error (*MAE*, kcal mol^*−*1^). The XGBoost model achieves an *R*^2^ of 0.92, a *RMSE* of 0.537 kcal mol^*−*1^, and an *MAE* of 0.411 kcal mol^*−*1^. The data points cluster tightly and uniformly along the y = x diagonal across the full range of experimental ΔG values, confirming that the trained model has successfully learned the mapping between the structural and thermodynamic feature space and the target binding affinity.

**Table 1:** Model evaluation matrix parameters.

| Model | Training Set |  |  | Test Set |  |  |
| --- | --- | --- | --- | --- | --- | --- |
| | $R^2$ | $RMSE$ | $MAE$ | $R^2$ | $RMSE$ | $MAE$ |
| XGBoost | 0.920 | 0.537 | 0.411 | 0.658 | 1.107 | 0.857 |
| Random Forest | 0.777 | 0.897 | 0.717 | 0.651 | 1.118 | 0.880 |
| Support Vector Machine | 0.841 | 0.757 | 0.517 | 0.613 | 1.177 | 0.922 |
| Decision Tree | 0.877 | 0.663 | 0.500 | 0.479 | 1.364 | 1.092 |

**Figure 3:**
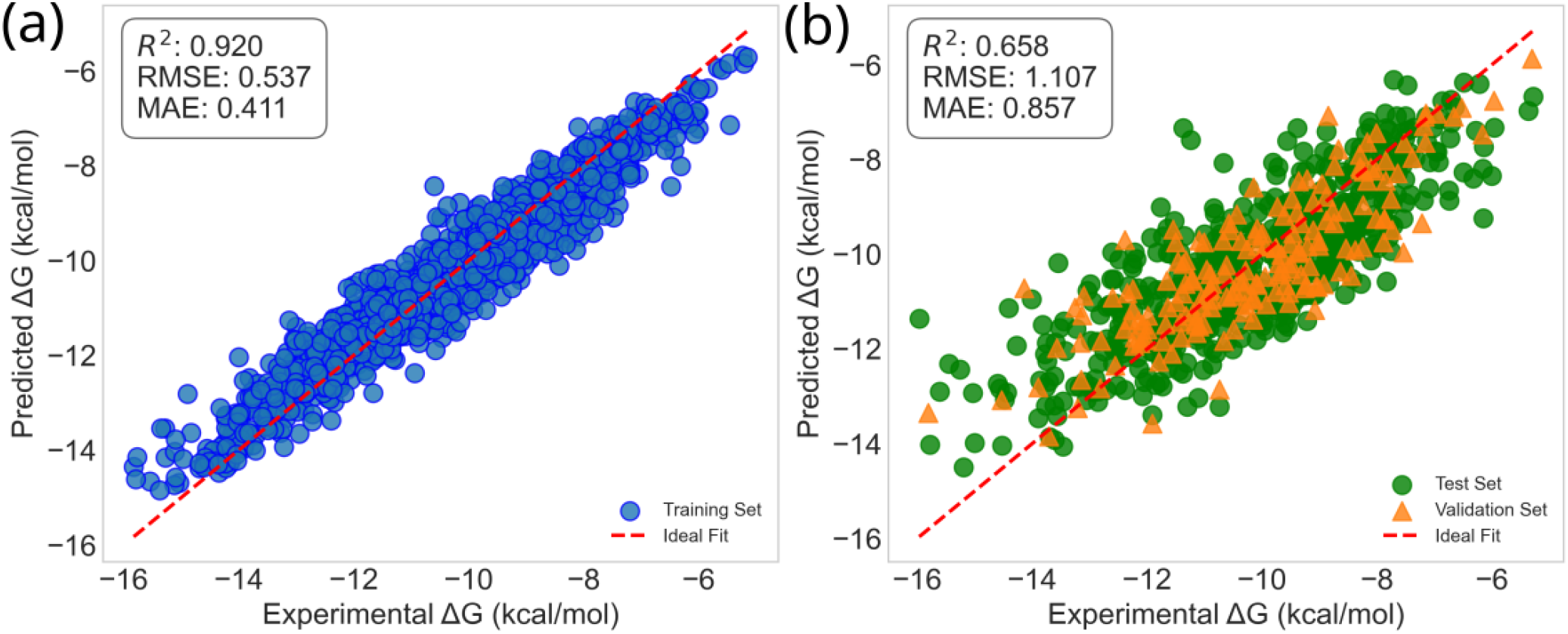
XGBoost-predicted vs experimentally determined protein-protein binding free energies (ΔG) for (a) the training set and (b) the combined test set and validation set. The dashed red diagonal line in each panel represents the locus of ideal agreement (y = x). Experimental ΔG values span the range -16 to -6 kcal mol^*−*1^, corresponding to the protein-protein binding affinities retained after outlier removal.

On the held-out test set and validation set (see Figure 3(b), the model yields an *R*^2^ of 0.658, an *RMSE* of 1.107 kcal mol^*−*1^, and an *MAE* of 0.857 kcal mol^*−*1^. The substantial reduction in *R*^2^ relative to the training set (Δ*R*^2^ = 0.262) indicates a degree of overfitting, in which the model has partially memorised training-set patterns that do not fully generalise to unseen complexes. Despite this, an *R*^2^ of 0.658 on the test set represents a meaningful level of generalisation for a structural-descriptor-based regression model applied to a heterogeneous dataset of protein–protein complexes, and compares favourably with benchmark values reported in the recent literature for analogous tasks (*R*^2^ = 0.40-0.65).^77^

Figure 3(b) shows key features of the model’s behaviour on unseen data. In the central affinity range (ΔG ≈ -12 to -8 kcal mol^*−*1^), both the test (green) and validation (orange) data points are roughly centred on the y = x line, indicating unbiased prediction in the most densely sampled part of the affinity distribution. However, at the extremes of the experimental range, a regression-to-the-mean effect appears: complexes with very strong binding (ΔG *<* -13 kcal mol^*−*1^) are consistently under-predicted, while weakly binding complexes (ΔG *>* -7 kcal mol^*−*1^) tend to be overpredicted. This compression of the predicted dynamic range is a common result of regularised ensemble methods applied to training distributions that undersample the tails of the target variable. Importantly, the validation set (orange triangles) shows a scatter pattern largely similar to that of the test set (green circles), confirming that the 10% validation split used for hyperparameter tuning did not create an overly optimistic bias in the reported test performance.

### Feature-to-Feature Correlation

To assess the degree of linear dependence among the 32 computed structural and energetic descriptors, a pairwise Pearson correlation matrix was constructed for the full dataset (see Figure 4). The resulting heatmap reveals a pronounced block-diagonal structure, indicating that the feature set comprises several internally correlated groups rather than a collection of independent descriptors. This multicollinearity has direct implications for model interpretability and subsequent feature selection strategy. The first and most prominent correlation block encompasses the absolute buried surface area (BSA) descriptors BSA total, BSA CC, BSA AA, and BSA PP, which exhibit pairwise correlation coefficients exceeding r = 0.85. This result is expected because the total BSA is arithmetically decomposed into charged (CC), apolar (AA), and polar (PP) contributions; any two sub-terms that together account for a large fraction of the total will necessarily co-vary. The percentage-composition counterparts (%BSA CC, %BSA AA, %BSA PP) display lower, though still appreciable, mutual correlations (r = 0.4-0.7) and exhibit weaker correlation with absolute BSA.

**Figure 4:**
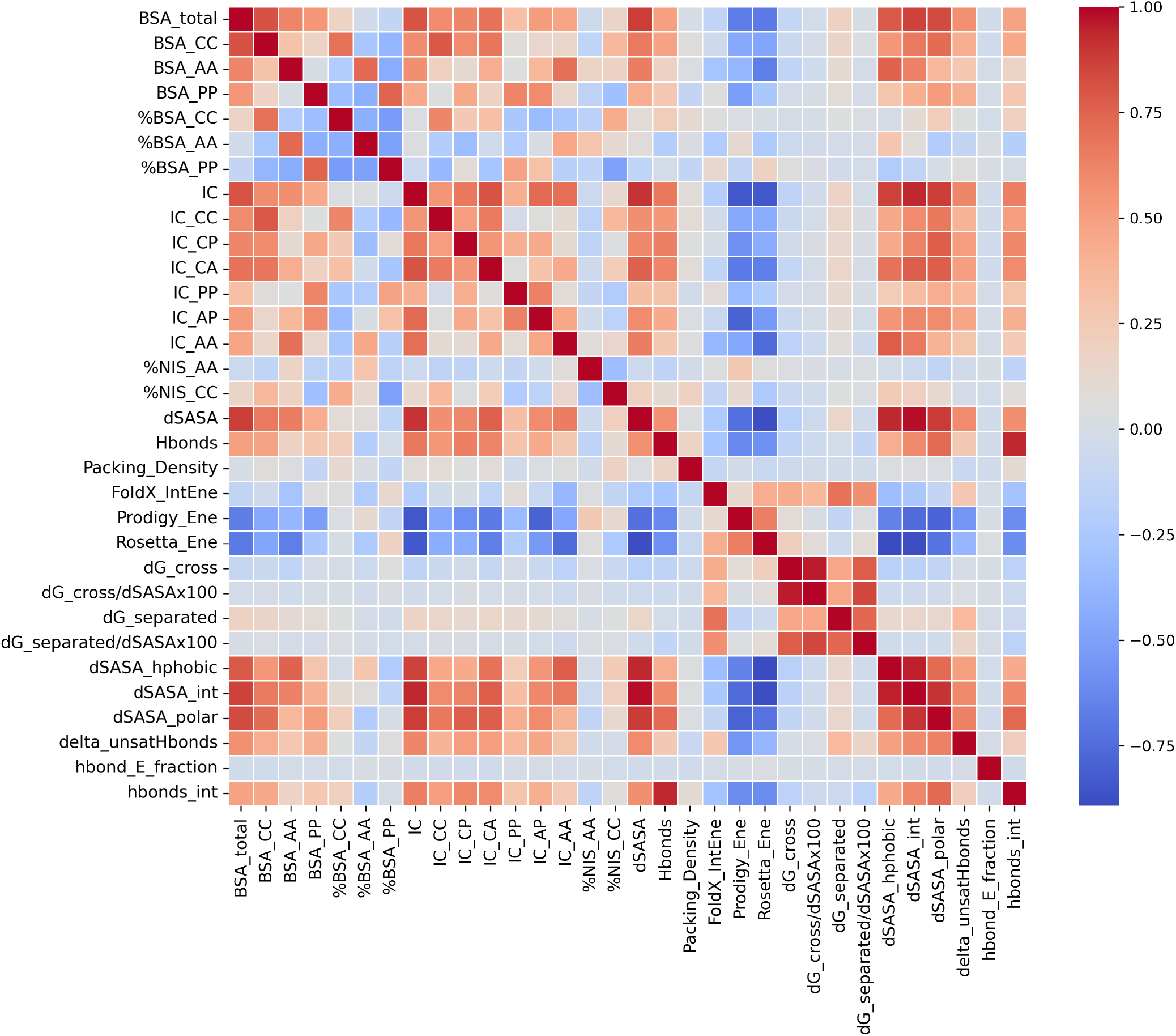
Feature-to-feature Pearson correlation matrix for the 32 structural and energetic descriptors computed for the protein-protein complex PDBbind+ dataset. Each cell reports the Pearson correlation coefficient between the corresponding feature pair. The color scale ranges from deep red (r = +1, perfect positive correlation) through white (r = 0, no linear correlation) to deep blue (r = −1, perfect negative correlation).

A second strongly correlated block comprises the interface contact (IC) descriptors, including total contacts (IC) and pair-resolved sub-types (IC CC, IC CP, IC CA, IC PP, IC AP, IC AA). Pairwise correlations within this group range from r = 0.65 to r *>* 0.90. Notably, IC is highly correlated with dSASA int and BSA total (r *>* 0.80), which is physically consistent: larger interfaces bury more surface area and simultaneously form more inter-residue contacts.

The change in solvent-accessible surface area sub-terms (dSASA int, dSASA hphobic, dSASA polar) form a tightly correlated cluster (r *>* 0.90 among sub-terms), as they are derived from the same parent quantity. These descriptors also show strong correlations with BSA total and IC, reinforcing the collinear relationship among geometric interface descriptors. In contrast, the non-interface surface composition features (%NIS AA, %NIS CC) are nearly orthogonal to both the BSA and IC blocks, exhibiting low cross-group correlations (r *<* 0.20)

Among the energetic descriptors, the four Rosetta-derived energy terms (d cross, dG cross/dSASAx100, dG separated, dG separated/dSASAx100) exhibit very strong intra-group correlations (r = 0.95 between raw and normalised variants of the same quantity), consistent with simple linear scaling by interface area. However, the cross-correlations between dG cross and dG separated are some-what lower (r = 0.75-0.85). Importantly, these energy terms show only moderate correlation with the geometric BSA and IC blocks (r = 0.30-0.55). The empirical and force-field-based energy scores (FoldX IntEne, Prodigy Ene, Rosetta Ene) show moderate mutual correlations (r = 0.50-0.70) and weak-to-moderate correlations with both the geometric and Rosetta energetic blocks.

The hydrogen-bond descriptors, hbonds int and hbond E fraction, form a tightly correlated pair (r *>* 0.90) but are largely independent of the geometric (BSA, IC) and energy blocks, with cross-group correlations generally below r = 0.35.

The delta unsatHbonds descriptor, which measures the energetic penalty associated with buried unsatisfied hydrogen-bond donors and acceptors, exhibits a weak negative correlation with hbonds int and negligible correlations with most geometric descriptors. In contrast, Packing Density shows only weak correlations with all descriptor classes (r *<* 0.40), indicating that steric complementarity is largely independent of interface size and interaction energetics. This descriptor’s relative independence from the dominant correlated clusters makes it a potentially valuable orthogonal feature in predictive models.

### Correlation between predicted and experimental ΔG of bNAbs across HIV-1 Clades

After extracting features from equilibrated 10-frames of the bNAb-HIV-1 complex structures for clades A, B, C, and G, the trained model, developed using the PDBbind^+^ database, was employed to predict the binding free energies (ΔG) for the HIV-1 datasets. All bNAbs used in the current study are listed in Table S1 (Supporting Information). The predicted ΔG (kcal mol^*−*1^) and the corresponding *K*_*D*_ (nM) of the bNAbs are shown in Tables S3 and S4 (Supporting Information), respectively.

To quantify the predicted ΔG against the reported experimental values for bNAbs, we plotted predicted and experimental data sets together in Figure 5 for HIV-1 clades A, B, C, and G. In panel 5(a), a moderate positive correlation (r = 0.367) is observed, indicating partial agreement between the predicted and experimental binding free energy.

**Figure 5:**
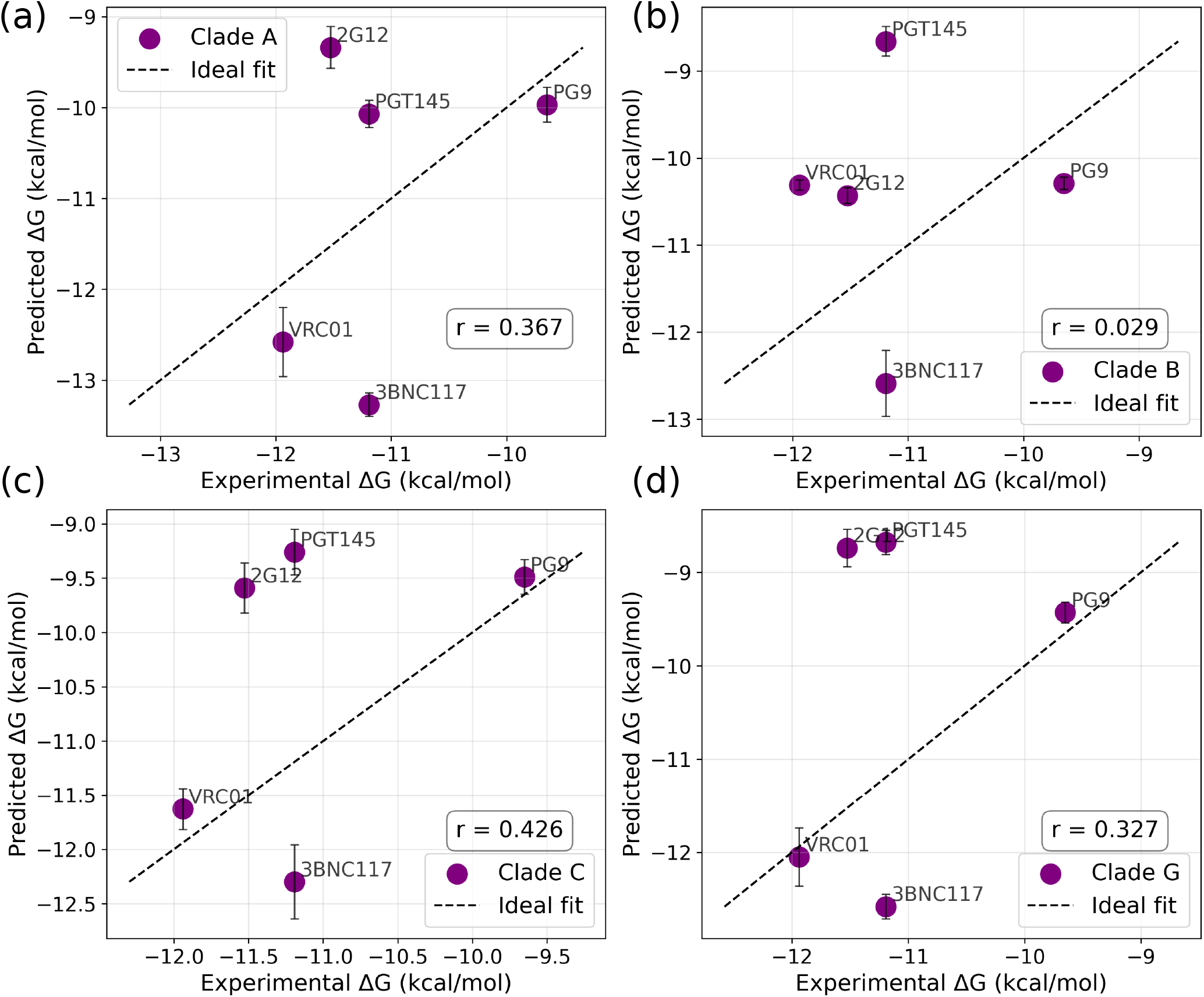
Scatter correlation plots of model predicted versus experimentally determined ΔG (kcal mol^*−*1^) of bNAbs across HIV-1 clades: (a) clade A, (b) clade B, (c) clade C, and (d) clade G. The standard deviation (STD) corresponds to the predicted ΔG of the extracted 10 to 11 different snapshots from last 1 ns of the simulation trajectory. The reported experimental binding affinities,^36^ (*K*_*D*_) were derived directly from binding energy Eq. 4.

The observed scatter reflects experimental complexity, including glycan heterogeneity and epitope accessibility, which are known to influence neutralization efficiency beyond static binding energetics. However, for clade B,Figure 5(b) shows weak correlation (r = 0.029) relative to other clades. Although potent antibodies, namely PG9, 2G12, VRC01, and 3BNC117, cluster toward more favorable predicted ΔG. The reduced correlation suggests that the structural and energetic features governing antibody-Env interactions in clade B are captured weekly by the predictive model.

In panel 5(c), a strong positive relationship between experimental and predicted ΔG values is revealed (r = 0.426) for clade C. While antibodies such as 2G12 and PGT145 maintain favorable predicted binding free energies, increased dispersion is evident compared to clades A and B. This behavior is consistent with experimental observations that clade C Envs exhibit greater glycan shielding and conformational variability, which can attenuate neutralization potency and flatten dose-response slopes despite strong intrinsic antibody affinity.^53^

Panel 5(d) showing a modest positive correlation (r = 0.327) for clade G. Potent antibodies such as PG9 and VRC01 display relatively consistent predicted binding affinities; however, deviations for antibodies like 3BNC117 suggest clade-specific structural effects. Experimental studies indicate that for clade G isolates, variability in neutralization slopes can arise from differences in glycan composition and Env trimer stability, leading to reduced predictability based on IC50 or ΔG alone.^53^

Across all clades, the analysis demonstrates that predicted binding free energy correlates moderately with experimentally derived ΔG values, but the strength of this relationship is clade dependent. Consistent with experimental literature, discrepancies arise because neutralization potency is governed not only by binding affinity but also by dose-response slope, which reflects cooperative and mechanistic aspects of antibody-virus interactions that are critical at therapeutic antibody concentrations.^53^

### Comparison of Model Predicted ΔG Across Clades

Figure 6 compares the epitope-wise ΔG of clinically evaluated bNAbs across different HIV-1 clades. CD4 binding sites (CD4bs) epitope (see cyan color region in Figure 1) target bNAbs show significant variation across clades. A close similarity in binding patterns is observed for several CD4bs targeting bNAbs, including 2411a, 12A21, BG24, N49P6, VRC-CH31, VRC-PG04, VRC-PG20, and VRC06b, as illustrated in Figure 6(a). In contrast, several bNAbs, including 3BNC117, HJ16, CH103, IOMA, NH49P7, VH1-46, and VRC01-FR3-03, exhibit distinct binding patterns across different HIV-1 clades. Among all antibodies analyzed, 3BNC117, b12, and N49P7 rank between those with the strongest binding affinities across all clades. Notably, 3BNC117 and N49P7 displays comparable ΔG for clades A but shows its weakest binding in clade C.

**Figure 6:**
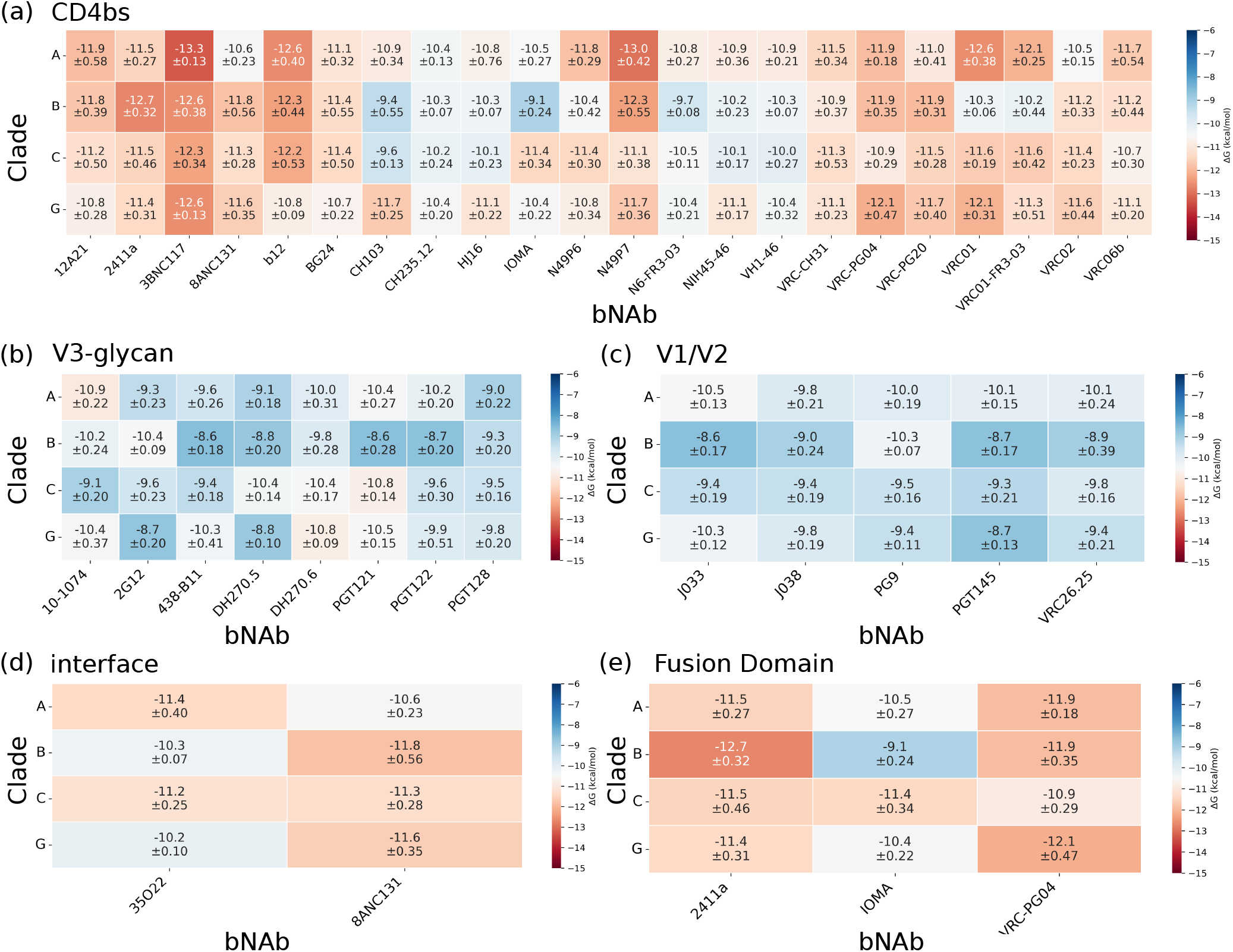
Clade-wise binding affinity of bNAbs comparison for the HIV-1 epitopes: (a) CD4bs, (b) V3-glycan, (c) V1V2 loop region, (d) gp120-gp41 Interface, and (e) Fusion Domain. The standard deviation (STD) has been calculated from the predicted ΔG of the extracted 10 to 11 different snapshots from the last 1 ns of the simulation trajectory.

In Figure 6(b), binding energy of V3-glycan (see blue color region in Figure 1) targeted bNAbs are compared across clades. The bNAbs DH270.5 and DH270.6 exhibit stronger binding affinities toward clade C compared to other clades. A similar clade-specific trend is observed for 10-1074, which shows enhanced binding in clade A. These observations suggest that these bNAbs are particularly effective in recognizing and binding the V3-glycan epitope regions within their respective clades. The bNAb 2G12 exhibits strong binding affinities toward clades B and C, while showing moderate binding in clades A and G, in quantitative agreement with experimental observations.^78–80^ In contrast, weaker binding is observed for bNAbs 438-B11 and PGT122 in clade B relative to other clades. Notably, PGT122 displays comparable ΔG for clades A and G.

The binding energies of bNAbs targeting the V1V2 epitopes (loop region) (see Figure 1) are shown in Figure 6(c). All bNAbs are showing the stronger affinity towards clade A, while bNAbs PG9 exhibit the higher binding affinities toward clade B with the moderate binding for clades C and G. This clade-dependent trend suggests that these bNAbs may exhibit a preferential specificity toward clade A relative to other clades. The bNAb J033 exhibits higher binding affinity toward clades A and G, with the weakest binding observed for clade B and moderate affinity for clade C. In contrast, the bNAb PGT145 shows strong binding affinities for clades A and C, while displaying the lowest binding strength for clade B and G.

Figure 6(d,e) illustrates the binding energies of bNAbs that target the gp120–gp41 interface and the fusion domain region (see Figure 1 for epitope clarity) of the HIV-1 Env trimer. As shown in Figure 6(d), the bNAb 35O22 exhibits high neutralization potential against the gp120-gp41 interface region, as reflected by its binding free energy values across different clades.^81,81^

Specifically, 35O22 demonstrates stronger binding affinities for clades A and C, moderate affinity for clades B and G. However, the bNAb 8ANC131 shows higher affinity for clade B compared to other clades. In Figure 6(e), the bNAb 2411a exhibits higher binding affinity toward clades B, C, and G, with moderate affinities observed for the remaining clades. Similarly, IOMA shows its strongest binding to clade C and the weakest affinity for clade B. In contrast, the bNAb VRC-PG04 displays enhanced binding affinity for clades A, B, and G, while maintaining moderate affinity across clade C.

### Variation in Predicted ΔG Across HIV-1 bNAb Epitope Classes

Figure 7 presents box-plot distributions of the predicted binding energies (ΔG, kcal mol^*−*1^) stratified by HIV-1 clade (A, B, C, and G) for five bNAbs epitope classes on the HIV-1 envelope glycoprotein (Env): (a) CD4 binding site (CD4bs), (b) V3-glycan, (c) V1V2-loop, (d) gp120-gp41 interface, and (e) fusion domain. Across all five sub-figures, predicted ΔG values fall within -13 to -8 kcal mol^*−*1^, consistent with the experimentally observed affinities of established bNAbs against these conserved Env epitopes.^82–84^ The inter-clade and inter-epitope variation in the predicted distributions provides quantitative structural evidence for the differential susceptibility of distinct HIV-1 clades to bNAb-mediated neutralisation, which is of direct relevance to the design of cross-clade HIV-1 vaccine immunogens.^24^

**Figure 7:**
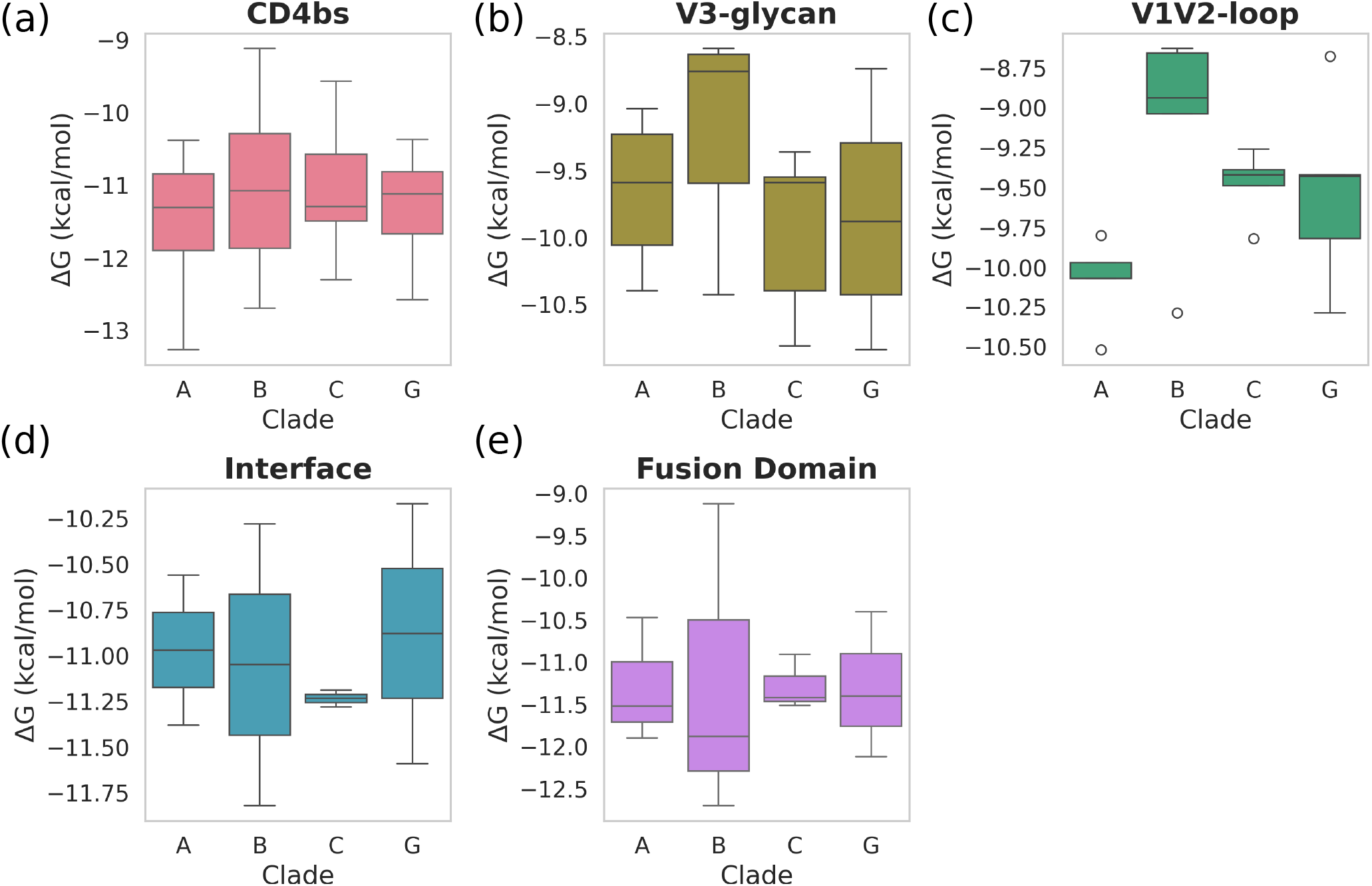
Box-plot distributions of XGBoost-predicted binding free energies (ΔG, kcal mol^*−*1^) for HIV-1 bNAb-Env complexes, stratified by clade (A, B, C, and G) and epitope class: (a) CD4bs, (b) V3-glycan, (c) V1V2-loop, (d) gp120-gp41 interface, and (e) fusion domain.

#### CD4 Binding Site (CD4bs)

Figure 7(a) shows the CD4bs panel tightly grouped median ΔG values across all four clades (range approximately -11.2 to -10.8 kcal mol^*−*1^), indicating that the predicted ΔG for bNAb engagement at the CD4bs is broadly conserved across clades A, B, C, and G. This is physically consistent with the functional requirement for conservation of the CD4 receptor-binding residues on gp120, which constitute the core of the CD4bs epitope and are shared among all major HIV-1 clades.^85^ The IQRs are widest for clade B (−12.0 to -10.2 kcal mol^*−*1^), reflecting the higher sequence diversity of clade B isolates at the CD4bs periphery, including variation in the surrounding V5 and D loops that modulate the approach geometry of CD4bs antibodies such as VRC01 and 3BNC117.^86^ The whiskers extend to approximately -13 kcal mol^*−*1^ for clade B, consistent with the highaffinity CD4bs antibody binding affinities (*K*_*D*_ = 0.1-1 nM) reported by ITC for VRC01-class bNAbs against diverse Env trimers.^86,87^

#### V3-glycan

The V3-glycan panel reveals a striking clade Bspecific pattern: clade B displays the widest IQR (approximately -9.5 to -8.5 kcal mol^*−*1^, upper whisker reaching -8.5 kcal mol^*−*1^) and the weakest predicted median affinity among all clades, in marked contrast to clades A, C, and G which show tighter distributions centred near -9.5 kcal mol^*−*1^. This unexpected result is counterintuitive given that bNAbs such as PGT121, PGT128, and 10-1074 were originally isolated and characterised against clade B isolates; however, it is consistent with experimental evidence that clade B Env trimers display considerable V3 tip and base sequence heterogeneity, which can reduce the predicted affinities of structural-feature-based models relative to the average experimental binding seen against matched isolates.^32^

The V3-glycan supersite is defined by contacts with the conserved N332 gp120 N-linked glycan and surrounding GDIR protein motif.^41^ The narrow lower whisker extending to approximately -10.5 kcal mol^*−*1^ for clade C reflects a subset of clade C isolates with long V1 loops that partially restrict access to the N332 glycan, reducing effective antibody approach angles and thus the model-predicted interface complementarity.^24^

#### V1V2-Loop

The V1V2-loop panel (Figure 7(c)) shows the most unusual clade-dependent pattern among all five classes. Clade A displays a compressed distribution with a very narrow IQR (−10.0 to -9.95 kcal mol^*−*1^), with multiple outliers extending to -10.6 kcal mol^*−*1^, while clade B shows a broad distribution centred near -8.8 kcal mol^*−*1^ with a narrow IQR and outlier at - 8.75 kcal mol^*−*1^. Clades C and G show intermediate distributions in the -9.5 to -9.3 kcal mol^*−*1^ range.

The compressed clade A distribution reflects the use of the BG505 (clade A) SOSIP trimer as the primary structural reference for V1V2-apex models-BG505 is the most extensively characterised Env trimer and has well-defined quaternary V1V2 apex geometry that is accurately encoded in the structural features.^88^ The weaker and narrower clade B predicted affinities may reflect greater structural variability in the clade B V1V2 loops used in the training dataset: V1V2-directed bNAbs such as PG9, PG16, and PGT145 engage a quaternary epitope requiring precise co-presentation of the N160 gp120 glycan and strand C residues, a geometry that is less consistently captured across diverse clade B structures than in the well-ordered clade A BG505 reference.^89^

#### gp120-gp41 Interface

The interface panel (Figure 7(d)) shows the overall strongest predicted affinities of all five epitope classes, with all clade medians falling between -11.25 and -10.75 kcal mol^*−*1^. Clade C exhibits the most compressed IQR (approximately -11.3 to -11.2 kcal mol^*−*1^) and the strongest median affinity, while clade B displays the broadest distribution with whiskers spanning -11.6 to -10.3 kcal mol^*−*1^.

The high predicted affinities across all clades are consistent with the structural conservation of the gp120-gp41 subunit junction, which is contacted by bNAbs such as 8ANC195.^90^ The narrow IQRs for clades A and C reflect the buried and sequence-invariant nature of the core gp120-gp41 contact residues across diverse HIV-1 clades, as their integrity is required for correct spike assembly and fusogenic function.^90,91^ Clade G shows a slightly wider distribution with a higher median than clades A and C (ΔG median -10.8 kcal mol^*−*1^), potentially reflecting minor structural rearrangements at the interface in clade G isolates identified by cryo-EM studies.^91^

#### Fusion Domain

The fusion domain panel (Figure 7(e)) shows that clade B displays the widest IQR (approximately -12.3 to -10.5 kcal mol^*−*1^), a markedly lower (stronger) median affinity than clades A, C, and G, and a lower whisker reaching -12.5 kcal mol^*−*1^. Clades A, C, and G show tighter distributions in the -11.5 to -11.0 kcal mol^*−*1^ range. The stronger and more heterogeneous predicted affinities for clade B are consistent with the origin of the principal fusion domain (FP) bNAb VRC34.01, which was characterised against clade B Env trimers and achieves its binding through critical contacts with Ile515 of the AVGIGAVFL fusion peptide motif.^92^ The tighter and weaker distributions for clades C and G are attributable to the higher frequency of Ile515Thr and adjacent FP sequence polymorphisms in non-clade-B isolates, which disrupt the hydrophobic packing geometry required for VRC34.01 engagement.^15,92^

Across all five panels, the predicted ΔG hierarchy follows: interface (−11.1 kcal mol^*−*1^) *>* fusion domain (−11.0) *>* CD4bs (−11.0) *>* V1V2-loop (−9.5) *>* V3-glycan (−9.5). This ordering is broadly consistent with the experimentally measured affinities of benchmark bNAbs for each class: interface and CD4bs bNAbs typically achieve sub-nanomolar KD values corresponding to ΔG *∼* -11 to -12 kcal mol^*−*1^, while V3-glycan and V1V2-directed bNAbs show greater variability due to glycan occupancy and quaternary Env conformation dependencies.^41,83,84^ The consistent pattern of wider IQRs for clade B across CD4bs, V3-glycan, and fusion domain panels reflects the higher Env sequence diversity of global clade B isolates relative to clade A, and the consequent broader range of predicted antibody–Env interface geometries in the structural dataset.^93,94^

### Neutralization Metrics of Predicted ΔG across HIV-1 Clades

The dose response curve (DRC) slope (m) describes the steepness of the sigmoidal neutralisation curve and quantifies the cooperativity of antibody-Env engagement. A slope of m = 1 indicates a simple non-cooperative doseresponse, whereas m *>* 1 reflects positive cooperativity, implying that neutralisation efficiency increases disproportionately with antibody concentration above the IC50. This distinction is critical for therapeutic applications: at clinically relevant antibody concentrations (typically 10-100 *µ*g mL^*−*1^), the instantaneous inhibitory potential (IIP) is proportional to m × log10(IC50), such that even a modest increase in slope from 1.0 to 1.5 translates to approximately 10 × greater reduction in viral load at equivalent antibody doses.^53,95^ The landmark study by Webb et al. demonstrated that, independent of IC50, CD4bs and V3-glycan bNAbs exhibit slopes *>* 1, whereas V2-glycan and gp41 epitope bNAbs exhibit slopes *<* 1, making slope an epitope-class-specific predictor of in vivo therapeutic effectiveness.^53,95^

Figure 8(a) reveals positive correlation between DRC slope and predicted ΔG. bNAbs with more negative (stronger) ΔG values exhibit steeper DRC slopes, irrespective of clade. The data cluster into two functionally distinct regimes separated by the slope = 1 boundary. In the sub-cooperative region (slope *<* 1), PG9 (clade G, slope = 0.77, ΔG = -9.2 ± 0.11 kcal mol^*−*1^), 3BNC117 (clade G, slope = 0.63, ΔG = -12.58 ± 0.13 kcal mol^*−*1^), and 2G12 (clades B, slope = 0.98) is found, indicating that despite achieving high predicted binding affinities, these antibodies fail to exhibit cooperative neutralisation against clade G isolates. This is consistent with the known sensitivity of PG9 and related V2-glycan apex antibodies to the quaternary Env conformation and glycan heterogeneity, which limits cooperative stoichiometric engagement of the trimer.^89^ The clade G data points for 3BNC117 in the sub-cooperative region are particularly notable: despite the most negative predicted ΔG of all antibodies shown (−12.58 ± 0.13 kcal mol^*−*1^), the shallow slope implies that high thermodynamic affinity does not guarantee cooperative neutralisation against clade G, likely reflecting partial occupation of non-neutralising epitope conformers on the clade G Env trimer surface.^45^

**Figure 8:**
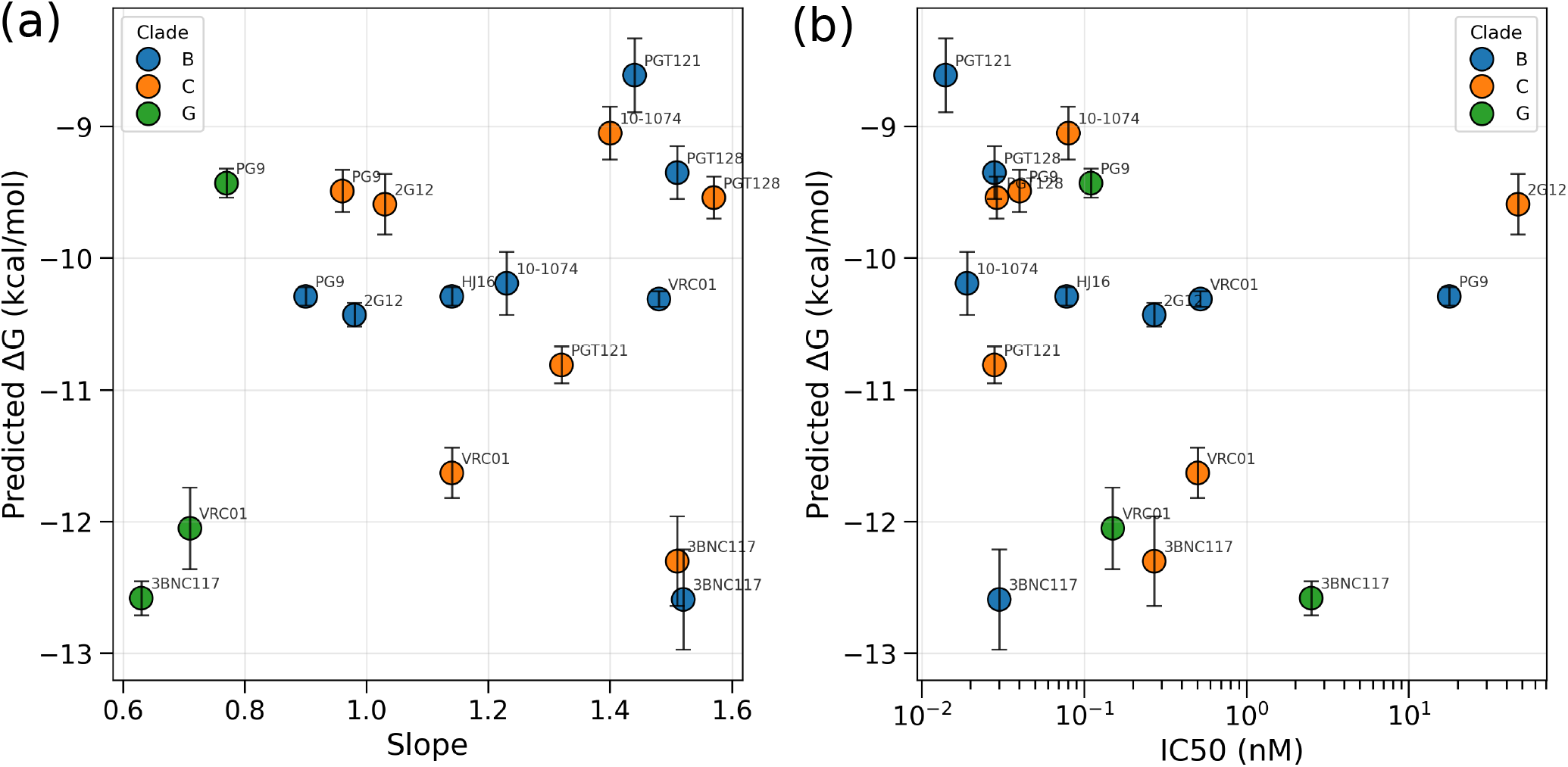
Correlation of predicted ΔG, kcal mol^*−*1^ with experimentally derived in vitro neutralisation parameters for eight HIV-1 bNAbs across clades B (blue), C (orange), and G (green). (a) DRC slope (m) vs ΔG. Slope *<* 1 indicates sub-cooperative and slope ≥ 1 indicates cooperative. (b) IC50 (nM, log scale) vs ΔG. IC50 *>* 1 nM indicates reduced potency, while IC50 ≤ indicates 1 nM high potency.

In the cooperative region (slope ≥ 1), bNAbs against the V3-glycan epitope-PGT121 (clade B, slope = 1.41, ΔG = -8.61 ± 0.28 kcal mol^*−*1^), PGT128 (clades B and C, slope = 1.51-1.57), and 10-1074 (clades B and C, slope = 1.23-1.4) cluster together with steep slopes, consistent with the published mean slopes of 1.59 (3BNC117 clade B) and 2 (PGT121) reported by Webb et al.^53^ CD4bs antibodies VRC01 (clades B and C, slope = 1.1-1.5) and HJ16 (clade B, slope = 1.14) also fall within the co-operative region, in agreement with the broad class-level observation that CD4bs bNAbs reliably achieve slopes *>* 1 against diverse Env panels.^53^ Across the cooperative region, more negative predicted ΔG values are associated with steeper slopes, suggesting that tighter thermodynamic binding promotes greater cooperativity, consistent with a model in which highaffinity occupancy of one Env protomer facilitates conformational recruitment of adjacent protomers in the pre-fusion trimer.^96^

Figure 8 (b) plots IC50 (nM, log scale) against predicted ΔG, revealing a negative correlation: more negative ΔG generally corresponds to lower (more potent) IC50. The IC50 = 1 nM boundary again separates the data into two regimes: antibodies with IC50 *<* 1 nM (green region, high potency) and IC50 *>* 1 nM (pink region, reduced potency). This thermodynamic-potency relationship is physically expected from the standard binding-neutralisation framework, wherein binding free energy determines antibody-Env occupancy probability at a given concentration, which in turn governs the fraction of Env trimers blocked per virion.^97^

In the high-potency regime (IC50 *<* 1 nM), PGT121 (clade B, IC50 0.014 nM, ΔG = -8.61 ± 0.28 kcal mol^*−*1^), PGT128 (clade B, IC50 = 0.028 nM), PG9 (clade B, IC50 = 17.7 nM), and 10-1074 (clade B, IC50 = 0.019 nM) are the most potent antibodies against clade B, consistent with published neutralisation panels reporting that V3-glycan bNAbs such as PGT121 achieve *∼*10-fold higher median potency than VRC01 against matched isolates.^32^ 3BNC117 (clade B and C, IC50 = 0.03-0.27 nM) and VRC01 (clade B, IC50 = 0.52 nM) also achieve sub-nanomolar IC50 against clade B, consistent with their well-characterised potency against tier-2 clade B reference viruses.^98^ HJ16 (clade B, IC50 = 0.078 nM, ΔG = -10.29 ± 0.07 kcal mol^*−*1^) shows strong predicted affinity and sub-nanomolar potency, consistent with its reported potency against a subset of CD4bs-sensitive isolates.^99^

Notably, several bNAbs, including VRC01 and 3BNC117, fall within the unfavorable region for clade G, exhibiting slopes of 0.65-0.70 (IC50 *∼* 2-10 nM), substantially below the critical threshold of 1, despite only modest differences in predicted ΔG relative to other clades. This observation highlights a key limitation of ΔG-based assessments, as small variations in binding free energy can translate into pronounced differences in neutralization efficacy. Given the global prevalence of clade G, these findings suggest that VRC01 and 3BNC117 may exhibit reduced therapeutic effectiveness as monotherapies in clade G infections, emphasizing that binding free energy alone may not fully capture in vivo neutralization potency.^53,86,100^

PG9, a V1V2-apex bNAb that recognizes the glycan-dependent quaternary epitope centred on N160-gp120,^101^ represents a notable outlier. Across clades B, C, and G, PG9 exhibits moderate binding affinities (ΔG ≈ -9.2 to -10.0 kcal mol^*−*1^) but consistently displays sub-unity slopes (0.70-0.85), indicative of incomplete neutralization. This behavior is consistent with the known sensitivity of PG9 to N160 glycan occupancy and V1V2 conformational heterogeneity, which can generate resistant viral subpopulations despite favorable binding energetics.^102,103^ Notably, the clade G PG9 variant exhibits a low IC50 (*∼* 0.07 nM) yet remains only partially effective because its slope remains below unity, emphasizing that strong binding affinity or low IC50 alone does not guarantee complete neutralization efficacy.^53^

The V3-glycan bNAbs PGT121, PGT128, and 10-1074 predominantly cluster within the favorable region for clades B and C, exhibiting slopes of 1.1-1.5, consistent with the characteristic steep neutralization profiles of this antibody class. For example, clade C 10-1074 and PGT121 display favorable slopes despite differences in predicted ΔG, and both retain potent IC50 values (≤ 0.2 nM). This agreement between structural and functional metrics supports the utility of ΔG as a predictor of neutralization potency for V3-glycan bNAbs against clades B and C. However, the absence of PGT121 and PGT128 activity against many clade G isolates, likely due to reduced N332 glycan occupancy, underscores a limitation of ΔG-based predictions, as glycan heterogeneity can substantially influence neutralization outcomes independent of binding energetics.^41^

It is clear that bNAbs VRC01, 3BNC117, PGT121, and PGT128 perform well to excellently in clade B, and have moderate to poor performance in clades C and G, respectively.

## Conclusion

In this study, an XGBoost machine learning model was trained using the PDBBind+ protein-protein complex database and demonstrated superior predictive performance compared to other regression models, achieving a training *R*^2^ of 0.92 and a test *R*^2^ of 0.658. The optimized model was subsequently applied to HIV-1 datasets to predict the binding free energies of multiple broadly neutralizing antibodies (bNAbs) across clades A, B, C, and G. Notably, clade B exhibits the largest variability in predicted binding affinities, spanning from approximately -8.5 to -13.0 kcal mol^*−*1^ across different epitopes.

The predicted binding free energies (ΔG) follow the trend: gp120-gp41 interface ≈ fusion domain *>* CD4 binding site (CD4bs) *>* V1V2 loop ≈ V3-glycan. This trend is broadly consistent with experimentally reported binding affinities of benchmark bNAbs within each epitope class. Antibodies targeting the gp120-gp41 interface and fusion domain typically exhibit sub-nanomolar dissociation constants (*K*_*D*_), corresponding to ΔG values of approximately -11 to -13 kcal mol^*−*1^. In contrast, V3-glycan and V1V2-directed bNAbs display greater variability in binding affinities, likely arising from their dependence on glycan composition and sensitivity to quaternary epitope presentation.

A consistent clade-dependent pattern is visible in clade B antibody-Env pairs (steep slopes and low IC50), while clade G is disproportionately found in the sub-cooperative/reduced-potency regions. This is consistent with the epidemiological observation that clade B HIV-1 isolates are generally more susceptible to multiple bNAb classes than clade G isolates, partly due to lower glycan density and shorter variable loops in clade B Env. Clade C pairs occupy an intermediate position, with some antibodies (3BNC117, VRC01, 10-1074) retaining potent IC50 values, and others (PG9, 2G12) showing markedly reduced potency, consistent with the known susceptibility profile of clade C isolates in published global neutralisation panels.

The predicted ΔG correlates with both dose response curve (DRC) slope and IC50, and the relationship is modulated by clade and epitope class. Antibodies appearing in the cooperative/high-potency regime in panel (e.g., 3BNC117 clade B, PGT121 clade B) also appear in the sub-nanomolar IC50 regime, and vice versa. This concordance validates the physical rationale underlying our structural feature-based ΔG predictions: stronger predicted binding affinity (ΔG more negative) is associated with deeper inhibitory potency at low antibody concentrations (lower IC50) and with more cooperative dose–response behaviour (higher slope), both of which predict greater in vivo antiviral efficacy. However, the presence of clade G outliers with strongly negative ΔG but sub-cooperative slopes underscores the limitation of purely thermodynamic predictions in capturing the full complexity of neutralisation, which additionally depends on antibody access kinetics, Env conformational dynamics, and viral escape pathways.

Overall, these results provide valuable insights into the binding affinities of clinically evaluated bNAbs across different HIV-1 clades and contribute to a deeper understanding of clade-specific interactions, with potential implications for rational vaccine design.

## Supporting information

The additional data in the Supporting Information file support the results presented in the manuscript.

## Data and Software Availability

All Python scripts and analysis codes used in this work are publicly available at the GitHub repository: https://github.com/kumarGit24/Protein-Protein-Complex-Affinity-Prediction

## Conflicts of interest

There are no conflicts of interest to declare.

## Acknowledgments

We thank DST, India, for funding under joint Indo-Taiwan program in science and technology (project #2023/IN-TW/02). Y.K. also acknowledges the computational support from the PKM Lab, Department of Physics, IISc Bangaluru.

## TOC Graphic

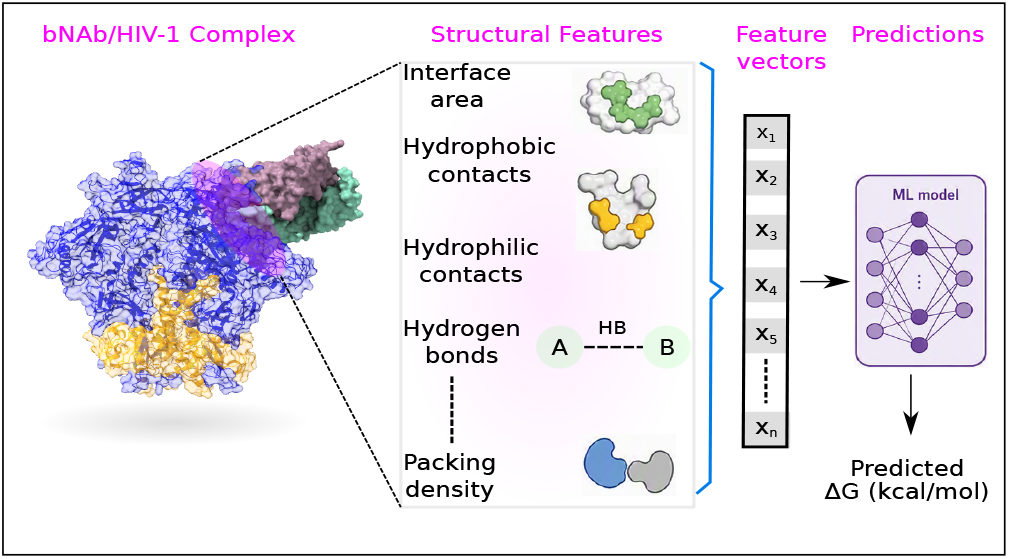

## Notes

### Competing Interest Statement

The authors have declared no competing interest.

## References

(1) Barré-Sinoussi, F.; Chermann, J.-C.; Rey, F.; Nugeyre, M. T.; Chamaret, S.; Gruest, J.; Dauguet, C.; Axler-Blin, C.; Vézinet-Brun, F.; Rouzioux, C.; others Isolation of a T-lymphotropic retrovirus from a patient at risk for acquired immune deficiency syndrome (AIDS). Science 1983, 220, 868–871.

(2) Caskey, M.; Klein, F.; Nussenzweig, M. C. Broadly neutralizing antibodies for HIV-1 prevention or immunotherapy. New England Journal of Medicine 2016, 375, 2019–2021.

(3) Wang, J.; Xiao, N.; Zhu, Z.; Qiao, H.; Zhao, F.; Zhang, L.; Gou, J.; Lu, M.; He, Y.; Lu, H.; others Comparing acute versus AIDS ART initiation on HIV-1 integration sites and clonal expansion. Signal Transduction and Targeted Therapy 2025, 10, 23.

(4) HIV data and statistics. 2025; https://www.who.int/teams/global-hiv-hepatitis-and-stis-programmes/hiv/strategic-information/hiv-data-and-statistics.

(5) Global HIV % AIDS statistics — Fact sheet. 2025; https://www.unaids.org/en/resources/fact-sheet.

(6) Tan, K.; Liu, J.-h.; Wang, J.-h.; Shen, S.; Lu, M. Atomic structure of a thermostable subdomain of HIV-1 gp41. Proceedings of the National Academy of Sciences 1997, 94, 12303–12308.

(7) Weissenhorn, W.; Dessen, A.; Harrison, S.; Skehel, J.; Wiley, D. Atomic structure of the ectodomain from HIV-1 gp41. Nature 1997, 387, 426–430.

(8) Wyatt, R.; Sodroski, J. The HIV-1 envelope glycoproteins: fusogens, antigens, and immunogens. Science 1998, 280, 1884–1888.

(9) Bazan, H. A.; Alkhatib, G.; Broder, C. C.; Berger, E. A. Patterns of CCR5, CXCR4, and CCR3 usage by envelope glycoproteins from human immunodeficiency virus type 1 primary isolates. Journal of virology 1998, 72, 4485–4491.

(10) Björndal, A.; Deng, H.; Jansson, M.; Fiore, J. R.; Colognesi, C.; Karlsson, A.; Albert, J.; Scarlatti, G.; Littman, D. R.; Fenyö, E. M. Coreceptor usage of primary human immunodeficiency virus type 1 isolates varies according to biological phenotype. Journal of virology 1997, 71, 7478–7487.

(11) Kowalski, M.; Potz, J.; Basiripour, L.; Dorfman, T.; Goh, W. C.; Terwilliger, E.; Dayton, A.; Rosen, C.; Haseltine, W.; Sodroski, J. Functional regions of the envelope glycoprotein of human immunodeficiency virus type 1. Science 1987, 237, 1351–1355.

(12) Lu, M.; Blacklow, S. C.; Kim, P. S. A trimeric structural domain of the HIV1 transmembrane glycoprotein. Nature structural biology 1995, 2, 1075–1082.

(13) Merk, A.; Subramaniam, S. HIV-1 envelope glycoprotein structure. Current opinion in structural biology 2013, 23, 268–276.

(14) Ward, A. B.; Wilson, I. A. Insights into the trimeric HIV-1 envelope glycoprotein structure. Trends in biochemical sciences 2015, 40, 101–107.

(15) Chen, B. Molecular mechanism of HIV-1 entry. Trends in microbiology 2019, 27, 878–891.

(16) Gorai, B.; Sahoo, A. K.; Srivastava, A.; Dixit, N. M.; Maiti, P. K. Concerted Interactions between Multiple gp41 Trimers and the Target Cell Lipidome May Be Required for HIV-1 Entry. Journal of Chemical Information and Modeling 2021, 61, 444–454.

(17) Freed, E. O.; Martin, M. A. The role of human immunodeficiency virus type 1 envelope glycoproteins in virus infection. Journal of Biological Chemistry 1995, 270, 23883–23886.

(18) Lataillade, M.; Lalezari, J. P.; Kozal, M.; Aberg, J. A.; Pialoux, G.; Cahn, P.; Thompson, M.; Molina, J.-M.; Moreno, S.; Grinsztejn, B.; others Safety and efficacy of the HIV-1 attachment inhibitor prodrug fostemsavir in heavily treatment-experienced individuals: week 96 results of the phase 3 BRIGHTE study. The Lancet HIV 2020, 7, e740–e751.

(19) Solomon, D. A.; Sax, P. E. Current state and limitations of daily oral therapy for treatment. Current Opinion in HIV and AIDS 2015, 10, 219–225.

(20) Archin, N. M.; Sung, J. M.; Garrido, C.; Soriano-Sarabia, N.; Margolis, D. M. Eradicating HIV-1 infection: seeking to clear a persistent pathogen. Nature Reviews Microbiology 2014, 12, 750–764.

(21) Kwong, P. D.; Mascola, J. R. Human antibodies that neutralize HIV-1: identification, structures, and B cell ontogenies. Immunity 2012, 37, 412–425.

(22) Kwong, P. D.; Mascola, J. R.; Nabel, G. J. Broadly neutralizing antibodies and the search for an HIV-1 vaccine: the end of the beginning. Nature Reviews Immunology 2013, 13, 693–701.

(23) Desikan, R.; Raja, R.; Dixit, N. M. Early exposure to broadly neutralizing antibodies may trigger a dynamical switch from progressive disease to lasting control of SHIV infection. PLoS computational biology 2020, 16, e1008064.

(24) Arandhara, M.; Kumar, Y.; Dixit, N. M.; Maiti, P. K. Clade-specific influences of glycans on the interactions between HIV1 envelope and broadly neutralizing antibodies. Journal of Chemical Information and Modeling 2025, 65, 13296–13314.

(25) Jardine, J. G.; Sok, D.; Julien, J.-P.; Briney, B.; Sarkar, A.; Liang, C.-H.; Scherer, E. A.; Henry Dunand, C. J.; Adachi, Y.; Diwanji, D.; others Minimally mutated HIV-1 broadly neutralizing antibodies to guide reductionist vaccine design. PLoS pathogens 2016, 12, e1005815.

(26) Mahomed, S. Broadly neutralizing antibodies for HIV prevention: a comprehensive review and future perspectives. Clinical Microbiology Reviews 2024, 37, e00152–22.

(27) Wagh, K.; Bhattacharya, T.; Williamson, C.; Robles, A.; Bayne, M.; Garrity, J.; Rist, M.; Rademeyer, C.; Yoon, H.; Lapedes, A.; others Optimal combinations of broadly neutralizing antibodies for prevention and treatment of HIV-1 clade C infection. PLoS pathogens 2016, 12, e1005520.

(28) Stewart-Jones, G. B.; Soto, C.; Lemmin, T.; Chuang, G.-Y.; Druz, A.; Kong, R.; Thomas, P. V.; Wagh, K.; Zhou, T.; Behrens, A.-J.; others Trimeric HIV-1-Env structures define glycan shields from clades A, B, and G. Cell 2016, 165, 813–826.

(29) Leonard, C. K.; Spellman, M. W.; Riddle, L.; Harris, R. J.; Thomas, J. N.; Gregory, T. Assignment of intrachain disulfide bonds and characterization of potential glycosylation sites of the type 1 recombinant human immunodeficiency virus envelope glycoprotein (gp120) expressed in Chinese hamster ovary cells. Journal of Biological Chemistry 1990, 265, 10373–10382.

(30) Geyer, H.; Holschbach, C.; Hunsmann, G.; Schneider, J. Carbohydrates of human immunodeficiency virus. Structures of oligosaccharides linked to the envelope glycoprotein 120. Journal of Biological Chemistry 1988, 263, 11760–11767.

(31) Mizuochi, T.; Spellman, M.; Larkin, M.; Solomon, J.; Basa, L.; Feizi, T. Carbohydrate structures of the humanimmunodeficiency-virus (HIV) recombinant envelope glycoprotein gp120 produced in Chinese-hamster ovary cells. Biochemical Journal 1988, 254, 599–603.

(32) Mouquet, H.; Scharf, L.; Euler, Z.; Liu, Y.; Eden, C.; Scheid, J. F.; Halper-Stromberg, A.; Gnanapragasam, P. N.; Spencer, D. I.; Seaman, M. S.; others Complex-type N-glycan recognition by potent broadly neutralizing HIV antibodies. Proceedings of the National Academy of Sciences 2012, 109, E3268– E3277.

(33) Kong, L.; Lee, J. H.; Doores, K. J.; Murin, C. D.; Julien, J.-P.; McBride, R.; Liu, Y.; Marozsan, A.; Cupo, A.; Klasse, P.-J.; others Supersite of immune vulnerability on the glycosylated face of HIV-1 envelope glycoprotein gp120. Nature structural & molecular biology 2013, 20, 796–803.

(34) Klasse, P.; Sanders, R. W.; Ward, A. B.; Wilson, I. A.; Moore, J. P. The HIV-1 envelope glycoprotein: structure, function and interactions with neutralizing antibodies. Nature Reviews Microbiology 2025, 1–19.

(35) Moore, P. L.; Murphy, M. K.; Derdeyn, C. A. HIV glycans in infection and immunity ; Springer, 2013; pp 85–115.

(36) Basu, S.; Gohain, N.; Kim, J.; Trinh, H. V.; Choe, M.; Joyce, M. G.; Rao, M. Determination of binding affinity of antibodies to HIV-1 recombinant envelope glycoproteins, pseudoviruses, infectious molecular clones, and cell-expressed trimeric gp160 using microscale thermophoresis. Cells 2023, 13, 33.

(37) Bontempo, A.; Garcia, M. M.; Rivera, N.; Cayabyab, M. J. A systematic approach to HIV-1 vaccine immunogen selection. AIDS research and human retroviruses 2020, 36, 762–770.

(38) Liao, H.-X.; Bonsignori, M.; Alam, S. M.; McLellan, J. S.; Tomaras, G. D.; Moody, M. A.; Kozink, D. M.; Hwang, K.-K.; Chen, X.; Tsao, C.-Y.; others Vaccine induction of antibodies against a structurally heterogeneous site of immune pressure within HIV-1 envelope protein variable regions 1 and 2. Immunity 2013, 38, 176–186.

(39) Alam, S. M.; Liao, H.-X.; Tomaras, G. D.; Bonsignori, M.; Tsao, C.-Y.; Hwang, K.-K.; Chen, H.; Lloyd, K. E.; Bowman, C.; Sutherland, L.; others Antigenicity and immunogenicity of RV144 vaccine AIDSVAX clade E envelope immunogen is enhanced by a gp120 N-terminal deletion. Journal of virology 2013, 87, 1554–1568.

(40) Rerks-Ngarm, S.; Pitisuttithum, P.; Nitayaphan, S.; Kaewkungwal, J.; Chiu, J.; Paris, R.; Premsri, N.; Namwat, C.; De Souza, M.; Adams, E.; others Vaccination with ALVAC and AIDSVAX to prevent HIV-1 infection in Thailand. New England Journal of Medicine 2009, 361, 2209–2220.

(41) Garces, F.; Lee, J. H.; de Val, N.; de la Pena, A.T.; Kong, L.; Puchades, C.; Hua, Y.; Stanfield, R. L.; Burton, D. R.; Moore, J. P.; others Affinity maturation of a potent family of HIV antibodies is primarily focused on accommodating or avoiding glycans. Immunity 2015, 43, 1053–1063.

(42) Nandy, B.; Bindu, D. H.; Dixit, N. M.; Maiti, P. K. Simulations reveal that the HIV-1 gp120-CD4 complex dissociates via complex pathways and is a potential target of the polyamidoamine (PAMAM) dendrimer. The Journal of chemical physics 2013, 139.

(43) Nandy, B.; Saurabh, S.; Sahoo, A. K.; Dixit, N. M.; Maiti, P. K. The SPL7013 dendrimer destabilizes the HIV-1 gp120– CD4 complex. Nanoscale 2015, 7, 18628–18641.

(44) Rajendra, D.; Maroli, N.; Dixit, N. M.; Maiti, P. K. Molecular dynamics simulations show how antibodies may rescue HIV-1 mutants incapable of infecting host cells. Journal of Biomolecular Structure and Dynamics 2025, 43, 2982–2992.

(45) Feng, M.; Bell, D. R.; Kang, H.; Shao, Q.; Zhou, R. Exploration of HIV-1 fusion peptide–antibody VRC34. 01 binding reveals fundamental neutralization sites. Physical Chemistry Chemical Physics 2019, 21, 18569–18576.

(46) Moyo, T.; Kitchin, D.; Moore, P. L. Targeting the N332-supersite of the HIV-1 envelope for vaccine design. Expert opinion on therapeutic targets 2020, 24, 499–509.

(47) Garg, A. K.; Desikan, R.; Dixit, N. M. Preferential presentation of high-affinity immune complexes in germinal centers can explain how passive immunization improves the humoral response. Cell reports 2019, 29, 3946–3957.

(48) Saha, A.; Dixit, N. M. Pre-existing resistance in the latent reservoir can compromise VRC01 therapy during chronic HIV-1 infection. PLOS Computational Biology 2020, 16, e1008434.

(49) LaMont, C.; Otwinowski, J.; Vanshylla, K.; Gruell, H.; Klein, F.; Nourmohammad, A. Design of an optimal combination therapy with broadly neutralizing antibodies to suppress HIV-1. Elife 2022, 11, e76004.

(50) Gorai, B.; Sahoo, A. K.; Srivastava, A.; Dixit, N. M.; Maiti, P. K. Concerted interactions between multiple gp41 trimers and the target cell lipidome may be required for HIV-1 entry. Journal of Chemical Information and Modeling 2020, 61, 444–454.

(51) Gorai, B.; Das, S.; Maiti, P. K. Prediction and validation of HIV-1 gp41 ecto-transmembrane domain post-fusion trimeric structure using molecular modeling. Journal of Biomolecular Structure and Dynamics 2020, 38, 2592–2603.

(52) Berman, H. M. The Protein Data Bank. Nucleic Acids Research 2000, 28, 235–242.

(53) Webb, N. E.; Montefiori, D. C.; Lee, B. Dose–response curve slope helps predict therapeutic potency and breadth of HIV broadly neutralizing antibodies. Nature communications 2015, 6, 8443.

(54) Los Alamos National Laboratory Neutralizing Antibody Contacts & Features Database. 2025; https://www.hiv.lanl.gov/content/immunology, Accessed: March 24, 2025.

(55) Wang, R.; Fang, X.; Lu, Y.; Wang, S. The PDBbind database: Collection of binding affinities for proteinligand complexes with known three-dimensional structures. Journal of medicinal chemistry 2004, 47, 2977–2980.

(56) Van Der Spoel, D.; Lindahl, E.; Hess, B.; Groenhof, G.; Mark, A. E.; Berendsen, H. J. GROMACS: fast, flexible, and free. Journal of computational chemistry 2005, 26, 1701–1718.

(57) Huang, J.; MacKerell Jr, A. D. CHARMM36 all-atom additive protein force field: Validation based on comparison to NMR data. Journal of computational chemistry 2013, 34, 2135–2145.

(58) Mark, P.; Nilsson, L. Structure and dynamics of the TIP3P, SPC, and SPC/E water models at 298 K. The Journal of Physical Chemistry A 2001, 105, 9954–9960.

(59) NOSÉ, S. U. I. A molecular dynamics method for simulations in the canonical ensemble. Molecular physics 2002, 100, 191–198.

(60) Hoover, W. G. Canonical dynamics: Equilibrium phase-space distributions. Physical review A 1985, 31, 1695.

(61) Parrinello, M.; Rahman, A. Polymorphic transitions in single crystals: A new molecular dynamics method. Journal of Applied physics 1981, 52, 7182–7190.

(62) Hess, B.; Bekker, H.; Berendsen, H. J.; Fraaije, J. G. LINCS: A linear constraint solver for molecular simulations. Journal of computational chemistry 1997, 18, 1463–1472.

(63) Darden, T.; York, D.; Pedersen, L.; others Particle mesh Ewald: An N log (N) method for Ewald sums in large systems. Journal of chemical physics 1993, 98, 10089–10089.

(64) Essmann, U.; Perera, L.; Berkowitz, M. L.; Darden, T.; Lee, H.; Pedersen, L. G. A smooth particle mesh Ewald method. The Journal of chemical physics 1995, 103, 8577–8593.

(65) Wang, R.; Fang, X.; Lu, Y.; Yang, C.Y.; Wang, S. The PDBbind database: methodologies and updates. Journal of medicinal chemistry 2005, 48, 4111–4119.

(66) Pettersen, E. F.; Goddard, T. D.; Huang, C. C.; Meng, E. C.; Couch, G. S.; Croll, T. I.; Morris, J. H.; Ferrin, T. E. UCSF ChimeraX: Structure visualization for researchers, educators, and developers. Protein science 2021, 30, 70–82.

(67) Michaud-Agrawal, N.; Denning, E. J.; Woolf, T. B.; Beckstein, O. MDAnalysis: a toolkit for the analysis of molecular dynamics simulations. Journal of computational chemistry 2011, 32, 2319–2327.

(68) McDonald, I. K.; Thornton, J. M. Satisfying hydrogen bonding potential in proteins. Journal of molecular biology 1994, 238, 777–793.

(69) Honorato, R. V.; Trellet, M. E.; Jiménez-García, B.; Schaarschmidt, J. J.; Giulini, M.; Reys, V.; Koukos, P. I.; Rodrigues, J. P.; Karaca, E.; van Zundert, G. C.; others The HADDOCK2. 4 web server for integrative modeling of biomolecular complexes. Nature protocols 2024, 19, 3219–3241.

(70) Xue, L. C.; Rodrigues, J. P.; Kastritis, P. L.; Bonvin, A. M.; Vangone, A. PRODIGY: a web server for predicting the binding affinity of protein–protein complexes. Bioinformatics 2016, 32, 3676–3678.

(71) Schymkowitz, J.; Borg, J.; Stricher, F.; Nys, R.; Rousseau, F.; Serrano, L. The FoldX web server: an online force field. Nucleic acids research 2005, 33, W382– W388.

(72) Alford, R. F.; Leaver-Fay, A.; Jeliazkov, J. R.; O’Meara, M. J.; DiMaio, F. P.; Park, H.; Shapovalov, M. V.; Renfrew, P. D.; Mulligan, V. K.; Kappel, K.; others The Rosetta all-atom energy function for macromolecular modeling and design. Journal of chemical theory and computation 2017, 13, 3031–3048.

(73) Chen, T.; Guestrin, C. Xgboost: A scalable tree boosting system. Proceedings of the 22nd acm sigkdd international conference on knowledge discovery and data mining. 2016; pp 785–794.

(74) Breiman, L. Random forests. Machine learning 2001, 45, 5–32.

(75) Cortes, C.; Vapnik, V. Support-vector networks. Machine learning 1995, 20, 273–297.

(76) Quinlan, J. R. Induction of decision trees. Machine learning 1986, 1, 81–106.

(77) Chaves, E. J.; Sartori, J.; Santos, W. M.; Cruz, C. H.; Mhrous, E. N.; Nacimento-Filho, M. F.; Ferraz, M. V.; Lins, R. D. Estimating absolute protein–protein binding free energies by a super learner model. Journal of chemical information and modeling 2025, 65, 2602–2609.

(78) Trkola, A.; Purtscher, M.; Muster, T.; Ballaun, C.; Buchacher, A.; Sullivan, N.; Srinivasan, K.; Sodroski, J.; Moore, J. P.; Katinger, H. Human monoclonal antibody 2G12 defines a distinctive neutralization epitope on the gp120 glycoprotein of human immunodeficiency virus type 1. Journal of virology 1996, 70, 1100–1108.

(79) West Jr, A. P.; Galimidi, R. P.; Foglesong, C. P.; Gnanapragasam, P. N.; Huey-Tubman, K. E.; Klein, J. S.; Suzuki, M. D.; Tiangco, N. E.; Vielmetter, J.; Bjorkman, P. J. Design and expression of a dimeric form of human immunodeficiency virus type 1 antibody 2G12 with increased neutralization potency. Journal of virology 2009, 83, 98–104.

(80) Wu, Y.; West, A. P.; Kim, H. J.; Thornton, M. E.; Ward, A. B.; Bjorkman, P. J. Structural basis for enhanced HIV-1 neutralization by a dimeric immunoglobulin G form of the glycan-recognizing antibody 2G12. Cell Reports 2013, 5, 1443–1455.

(81) Huang, J.; Kang, B. H.; Pancera, M.; Lee, J. H.; Tong, T.; Feng, Y.; Imamichi, H.; Georgiev, I. S.; Chuang, G.-Y.; Druz, A.; others Broad and potent HIV-1 neutralization by a human antibody that binds the gp41–gp120 interface. Nature 2014, 515, 138–142.

(82) Kwong, P. D.; Doyle, M. L.; Casper, D. J.; Cicala, C.; Leavitt, S. A.; Majeed, S.; Steenbeke, T. D.; Venturi, M.; Chaiken, I.; Fung, M.; others HIV-1 evades antibody-mediated neutralization through conformational masking of receptor-binding sites. Nature 2002, 420, 678–682.

(83) Kwong, P. D.; Mascola, J. R. HIV-1 vaccines based on antibody identification, B cell ontogeny, and epitope structure. Immunity 2018, 48, 855–871.

(84) Burton, D. R.; Mascola, J. R. Antibody responses to envelope glycoproteins in HIV-1 infection. Nature immunology 2015, 16, 571–576.

(85) Pancera, M.; Zhou, T.; Druz, A.; Georgiev, I. S.; Soto, C.; Gorman, J.; Huang, J.; Acharya, P.; Chuang, G.-Y.; Ofek, G.; others Structure and immune recognition of trimeric pre-fusion HIV-1 Env. Nature 2014, 514, 455–461.

(86) Wu, X.; Yang, Z.-Y.; Li, Y.; Hogerkorp, C.-M.; Schief, W. R.; Seaman, M. S.; Zhou, T.; Schmidt, S. D.; Wu, L.; Xu, L.; others Rational design of envelope identifies broadly neutralizing human monoclonal antibodies to HIV-1. Science 2010, 329, 856–861.

(87) Li, Y.; O’Dell, S.; Walker, L. M.; Wu, X.; Guenaga, J.; Feng, Y.; Schmidt, S. D.; McKee, K.; Louder, M. K.; Ledgerwood, J. E.; others Mechanism of neutralization by the broadly neutralizing HIV-1 monoclonal antibody VRC01. Journal of virology 2011, 85, 8954–8967.

(88) Lee, J. H.; Ozorowski, G.; Ward, A. B. Cryo-EM structure of a native, fully glycosylated, cleaved HIV-1 envelope trimer. Science 2016, 351, 1043–1048.

(89) Doria-Rose, N. A.; Schramm, C. A.; Gorman, J.; Moore, P. L.; Bhiman, J. N.; DeKosky, B. J.; Ernandes, M. J.; Georgiev, I. S.; Kim, H. J.; Pancera, M.; others Developmental pathway for potent V1V2-directed HIV-neutralizing antibodies. Nature 2014, 509, 55–62.

(90) Scharf, L.; Wang, H.; Gao, H.; Chen, S.; McDowall, A. W.; Bjorkman, P. J. Broadly neutralizing antibody 8ANC195 recognizes closed and open states of HIV1 Env. Cell 2015, 162, 1379–1390.

(91) Huang, J.; Ofek, G.; Laub, L.; Louder, M. K.; Doria-Rose, N. A.; Longo, N. S.; Imamichi, H.; Bailer, R. T.; Chakrabarti, B.; Sharma, S. K.; others Broad and potent neutralization of HIV-1 by a gp41-specific human antibody. Nature 2012, 491, 406–412.

(92) Kong, R.; Xu, K.; Zhou, T.; Acharya, P.; Lemmin, T.; Liu, K.; Ozorowski, G.; Soto, C.; Taft, J. D.; Bailer, R. T.; others Fusion peptide of HIV-1 as a site of vulnerability to neutralizing antibody. Science 2016, 352, 828–833.

(93) Gaschen, B.; Taylor, J.; Yusim, K.; Foley, B.; Gao, F.; Lang, D.; Novitsky, V.; Haynes, B.; Hahn, B. H.; Bhattacharya, T.; others Diversity considerations in HIV-1 vaccine selection. Science 2002, 296, 2354–2360.

(94) Sok, D.; Pauthner, M.; Briney, B.; Lee, J. H.; Saye-Francisco, K. L.; Hsueh, J.; Ramos, A.; Le, K. M.; Jones, M.; Jardine, J. G.; others A prominent site of antibody vulnerability on HIV envelope incorporates a motif associated with CCR5 binding and its camouflaging glycans. Immunity 2016, 45, 31–45.

(95) Shen, L.; Peterson, S.; Sedaghat, A. R.; McMahon, M. A.; Callender, M.; Zhang, H.; Zhou, Y.; Pitt, E.; Anderson, K. S.; Acosta, E. P.; others Dose-response curve slope sets class-specific limits on inhibitory potential of anti-HIV drugs. Nature medicine 2008, 14, 762–766.

(96) Julien, J.-P.; Cupo, A.; Sok, D.; Stanfield, R. L.; Lyumkis, D.; Deller, M. C.; Klasse, P.-J.; Burton, D. R.; Sanders, R. W.; Moore, J. P.; others Crystal structure of a soluble cleaved HIV-1 envelope trimer. Science 2013, 342, 1477–1483.

(97) Bonsignori, M.; Liao, H.-X.; Gao, F.; Williams, W. B.; Alam, S. M.; Montefiori, D. C.; Haynes, B. F. Antibody-virus co-evolution in HIV infection: paths for HIV vaccine development. Immunological reviews 2017, 275, 145–160.

(98) Mkhize, N. N.; Yssel, A. E.; Kaldine, H.; van Dorsten, R. T.; Woodward Davis, A. S.; Beaume, N.; Matten, D.; Lambson, B.; Modise, T.; Kgagudi, P.; others Neutralization profiles of HIV-1 viruses from the VRC01 Antibody Mediated Prevention (AMP) trials. PLoS pathogens 2023, 19, e1011469.

(99) Corti, D.; Langedijk, J. P.; Hinz, A.; Seaman, M. S.; Vanzetta, F.; Fernandez-Rodriguez, B. M.; Silacci, C.; Pinna, D.; Jarrossay, D.; Balla-Jhagjhoorsingh, S.; others Analysis of memory B cell responses and isolation of novel monoclonal antibodies with neutralizing breadth from HIV-1-infected individuals. PloS one 2010, 5, e8805.

(100) Corey, L.; Gilbert, P. B.; Juraska, M.; Montefiori, D. C.; Morris, L.; Karuna, S. T.; Edupuganti, S.; Mgodi, N. M.; Decamp, A. C.; Rudnicki, E.; others Two randomized trials of neutralizing antibodies to prevent HIV-1 acquisition. New England Journal of Medicine 2021, 384, 1003–1014.

(101) McLellan, J. S.; Pancera, M.; Carrico, C.; Gorman, J.; Julien, J.-P.; Khayat, R.; Louder, R.; Pejchal, R.; Sastry, M.; Dai, K.; others Structure of HIV-1 gp120 V1/V2 domain with broadly neutralizing antibody PG9. Nature 2011, 480, 336–343.

(102) McCoy, L. E.; Falkowska, E.; Doores, K. J.; Le, K.; Sok, D.; van Gils, M. J.; Euler, Z.; Burger, J. A.; Seaman, M. S.; Sanders, R. W.; others Incomplete neutralization and deviation from sigmoidal neutralization curves for HIV broadly neutralizing monoclonal antibodies. PLoS pathogens 2015, 11, e1005110.

(103) Ringe, R. P.; Colin, P.; Ozorowski, G.; Allen, J. D.; Yasmeen, A.; Seabright, G. E.; Lee, J. H.; Antanasijevic, A.; Rantalainen, K.; Ketas, T.; others Glycan heterogeneity as a cause of the persistent fraction in HIV-1 neutralization. PLoS Pathogens 2023, 19, e1011601.

