## Supplementary material for "Decoding Clade-Specific bNAb–HIV-1 Env Interactions Using Machine Learning": The additional data in the Supporting Information file support the results presented in the manuscript.

### Supporting Information
### for
### Decoding Clade-Specific bNAb-HIV-1 Env Interactions Using Machine Learning

Yogendra Kumar,[†] Shiang-Tai Lin,[‡] Narendra M. Dixit,[¶] and Prabal K. Maiti[*,†]

†*Centre for Condensed Matter Theory, Department of Physics, Indian Institute of Science, Bengaluru 560012, India*

‡*Department of Chemical Engineering, National Taiwan University, Taipei 106319, Taiwan*

¶*Department of Chemical Engineering, Indian Institute of Science, Bengaluru 560012, India*

§*Department of Bioengineering, Indian Institute of Science, Bengaluru 560012, India*

#### Contents

#### List of Figures

S1 Schematic for superimposition of RCSB obtained structure (orange) with missing AA modelled HIV-1 Env trimer (blue). In the final structure, the superimposed gp120 chains are deleted. . . . . . . . . . . . . . . . . . . . . . . S-3

S2 Experimental $\Delta$G distribution used in ML model training. The data range from -16 to -4 kcal mol$^{-1}$ was used for training, testing, and validation. . . . S-3

### List of Tables

S1 Epitope wise clinically trial bNAb listing used to predict the binding affinity or binding energy. We have a total of 30 CD4bs, seven V2-apex, nine V3-glycan class,two gp120/gp41 Interface, and one each of fusion peptide and glycan class bNAbs. . . . . . . . . . . . . . . . . . . . . . . . . . . . . . . S-4

S2 Summary of correlated feature groups identified from the Pearson correlation matrix. Representative features, their physical interpretation, and key correlation observations are listed for each group. . . . . . . . . . . . . . . . . S-5

S3 Predicted binding free energies ($\Delta G$, kcal mol$^{-1}$) of bNAbs across HIV-1 Env clades. Values are reported as mean ± STD. . . . . . . . . . . . . . . . . . . S-6

S4 Predicted dissociation constants ($K_d$, nM) of bNAbs across HIV-1 Env clades. Values are reported as mean ± STD. The expression $\Delta$G = RTln$K_d$ was used to get the dissociation constant of each bNAb, where R = 0.001987 kcal mol$^{-1}$ K$^{-1}$ and T = 300 K. . . . . . . . . . . . . . . . . . . . . . . . . . . . . . S-7

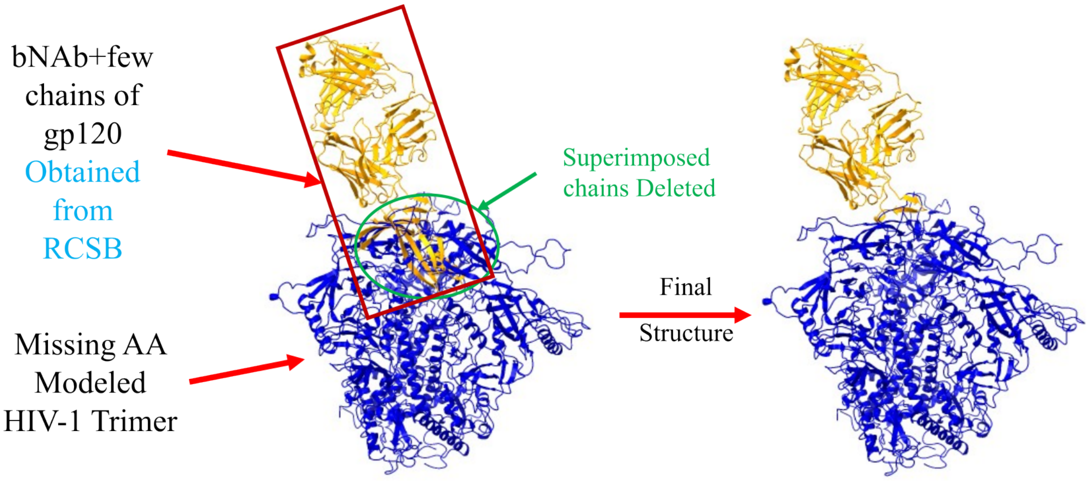


Figure S1: Schematic for superimposition of RCSB obtained structure (orange) with missing AA modelled HIV-1 Env trimer (blue). In the final structure, the superimposed gp120 chains are deleted.

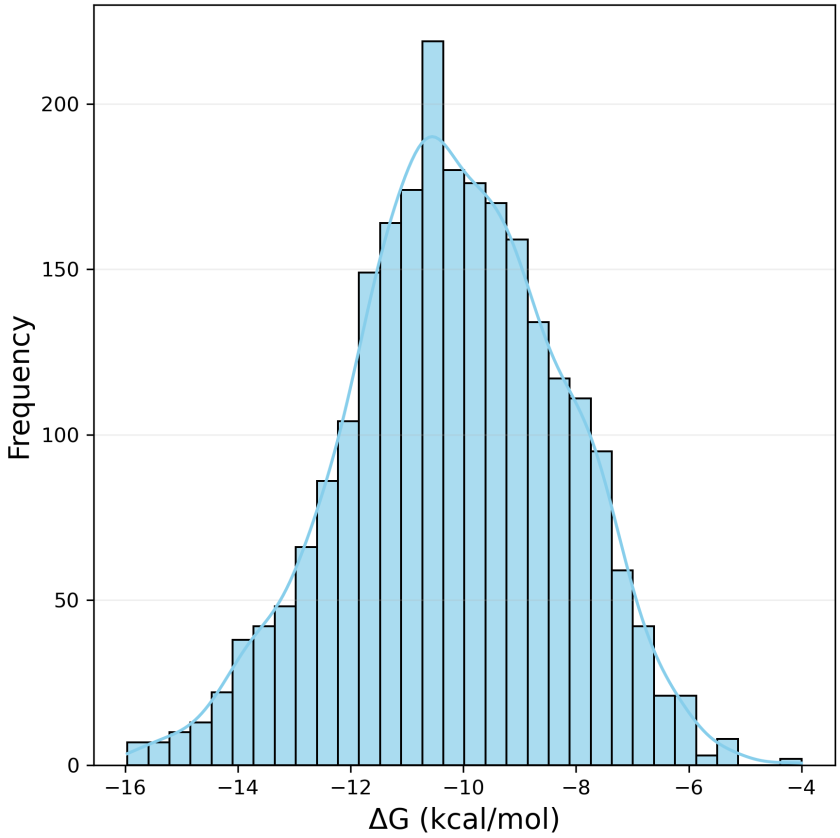


Figure S2: Experimental ΔG distribution used in ML model training. The data range from -16 to -4 kcal $mol^{-1}$ was used for training, testing, and validation.

Table S1: Epitope wise clinically trial bNAb listing used to predict the binding affinity or binding energy. We have a total of 30 CD4bs, seven V2-apex, nine V3-glycan class,two gp120/gp41 Interface, and one each of fusion peptide and glycan class bNAbs.

| bNAb | PDBID | Epitope | bNAb | PDBID | Epitope |
|---|---|---|---|---|---|
| VRC13 | 4YDJ | CD4bs | CD4 | 2QAD | CD4bs |
| BG24 | 7UCG | CD4bs | N6 | 5TE7 | CD4bs |
| VRC-PG04 | 3SE9 | CD4bs | b12 | 2NY7 | CD4bs |
| NIH45-46 | 4JDT | CD4bs | CH103 | 4JAN | CD4bs |
| 3BNC117 | 4JPV | CD4bs | HJ16 | 4YE4 | CD4bs |
| 12A21 | 4JPW | CD4bs | VRC26.25 | 6VTT | V2-apex |
| VRC-CH31 | 4LSP | CD4bs | J038 | 7MXD | V2-apex |
| VRC-PG20 | 4LSU | CD4bs | J033 | 7N28 | V2-apex |
| VRC07 | 4OLZ | CD4bs | PGT145 | 6NIJ | V2-apex |
| 8ANC131 | 4RWY | CD4bs | PG9 | 3U2S | V2-apex |
| VRC02 | 4RX4 | CD4bs | VRC38.01 | 5VGJ | V2-apex |
| VRC06b | 4XNZ | CD4bs | PG16 | 6ULC | V2-apex |
| 1B2530 | 4YFL | CD4bs | PGT135 | 4JM2 | V3-glycan |
| DRVIA7 | 5CD5 | CD4bs | DH270.6 | 6UM6 | V3-glycan |
| CH235.12 | 5F96 | CD4bs | 438-B11 | 6UTK | V3-glycan |
| CH235.9 | 5F9O | CD4bs | DH270.5 | 8SAS | V3-glycan |
| 3BNC55 | 5I9Q | CD4bs | PGT122 | 4TVP | V3-glycan |
| CAP257 | 5T33 | CD4bs | PGT121 | 5CEZ | V3-glycan |
| N60P23 | 5WB9 | CD4bs | 10-1074 | 7UCG | V3-glycan |
| VRC01 | 4XVT | CD4bs | BG18 | 6DFH | V3-glycan |
| N49P7 | 6BCK | CD4bs | PGT128 | 3TYG | V3-glycan |
| IOMA | 5T3Z | CD4bs | 35O22 | 4TVP | gp120/gp41 Interface |
| NC-Cow1 | 6OPA | CD4bs | 8ANC195 | 4P9H | gp120/gp41 Interface |
| N49P6 | 6OZ2 | CD4bs | PGT151 | 6DCQ | gp120/gp41 Interface |
| 2411a | 7JKS | CD4bs | 2G12 | 6OZC | glycan |
| 179NC75 | 7LLK | CD4bs | | | |

Table S2: Summary of correlated feature groups identified from the Pearson correlation matrix. Representative features, their physical interpretation, and key correlation observations are listed for each group.

| Feature Group | Physical Interpretation | Key Correlation |
|---|---|---|
| BSA_total, BSA_CC, BSA_AA, BSA_PP | Buried surface area (total, charged, apolar, polar) | Strong mutual correlations (r >0.85); size-dependent collinear block |
| %BSA_CC, %BSA_AA, %BSA_PP | Percentage composition of BSA | Moderate intra-group correlations; partially independent of absolute BSA |
| IC, IC_CC, IC_CP, IC_CA, IC_PP, IC_AP, IC_AA | Interface contacts (total and pair-type) | High internal correlations (r >0.80); IC highly correlated with dSASA |
| %NIS_AA, %NIS_CC | Non-interface surface composition | Near-orthogonal to BSA and IC blocks; low cross-group correlation |
| dSASA, dSASA_hphobic, dSASA_int, dSASA_polar | Change in SASA on binding | Strong correlation among dSASA sub-terms (r >0.90); collinear with BSA |
| dG_cross, dG_cross/dSASAx100, dG_separated, dG_separated/dSASAx100 | Rosetta binding free energies | Strong intra-pair correlations (r ∼0.95); partially decorrelated from geometric features |
| FoldX_IntEne, Prodigy_Ene, Rosetta_Ene | Empirical/FF-based energy scores | Moderate mutual correlations (r ∼0.5–0.7); weakly correlated with BSA/IC |
| hbond_E_fraction, hbonds_int | Hydrogen-bond count and energy fraction | Strong mutual correlation (r >0.90); near-independent of BSA/IC blocks |
| delta_unsatHbonds | Buried unsatisfied H-bonds penalty | Largely independent; weak negative correlation with hbonds_int |
| Packing_Density, packstat | Interface packing quality | Weakly correlated with geometric and energetic blocks |

Table S3: Predicted binding free energies ($\Delta G$, kcal mol$^{-1}$) of bNAbs across HIV-1 Env clades. Values are reported as mean ± STD.

| bNAb | Clade A | Clade B | Clade C | Clade G |
|---|---|---|---|---|
| 10-1074 | -10.94 ± 0.22 | -10.19 ± 0.24 | -9.05 ± 0.20 | -10.41 ± 0.37 |
| 12A21 | -11.89 ± 0.58 | -11.76 ± 0.39 | -11.25 ± 0.50 | -10.81 ± 0.28 |
| 179NC75 | -9.24 ± 0.22 | -9.16 ± 0.36 | -10.11 ± 0.10 | -8.79 ± 0.22 |
| 1B2530 | -10.90 ± 0.26 | -10.73 ± 0.34 | -10.65 ± 0.18 | -10.41 ± 0.18 |
| 2411a | -11.52 ± 0.27 | -12.70 ± 0.32 | -11.51 ± 0.46 | -11.40 ± 0.31 |
| 2G12 | -9.34 ± 0.23 | -10.43 ± 0.09 | -9.59 ± 0.23 | -8.78 ± 0.18 |
| 35O22 | -11.38 ± 0.40 | -10.28 ± 0.07 | -11.19 ± 0.25 | -10.17 ± 0.10 |
| 3BNC117 | -13.27 ± 0.13 | -12.59 ± 0.38 | -12.30 ± 0.34 | -12.58 ± 0.13 |
| 3BNC55 | -11.74 ± 0.34 | -12.15 ± 0.15 | -12.24 ± 0.23 | -11.85 ± 0.33 |
| 438-B11 | -9.59 ± 0.26 | -8.59 ± 0.18 | -9.36 ± 0.18 | -10.32 ± 0.41 |
| 8ANC131 | -10.56 ± 0.23 | -11.82 ± 0.56 | -11.28 ± 0.28 | -11.59 ± 0.35 |
| b12 | -12.64 ± 0.40 | -12.34 ± 0.44 | -12.22 ± 0.53 | -8.64 ± 0.00 |
| b13 | -14.04 ± 0.06 | -10.29 ± 0.07 | -10.62 ± 0.06 | -13.92 ± 0.08 |
| BG24 | -11.13 ± 0.32 | -11.37 ± 0.55 | -11.38 ± 0.50 | -10.71 ± 0.22 |
| CAP257-RH1 | -10.01 ± 0.26 | -9.63 ± 0.23 | -10.17 ± 0.27 | -9.47 ± 0.21 |
| CH103 | -10.89 ± 0.34 | -9.36 ± 0.55 | -9.57 ± 0.13 | -11.69 ± 0.25 |
| CH235.12 | -10.38 ± 0.13 | -10.29 ± 0.07 | -10.19 ± 0.24 | -10.40 ± 0.20 |
| CH235.9 | -10.71 ± 0.70 | -10.29 ± 0.07 | -10.62 ± 0.06 | -8.64 ± 0.00 |
| DH270.5 | -9.12 ± 0.18 | -8.76 ± 0.20 | -10.36 ± 0.14 | -8.84 ± 0.10 |
| DH270.6 | -9.96 ± 0.31 | -9.84 ± 0.28 | -10.44 ± 0.17 | -8.64 ± 0.00 |
| DRVIA7 | -11.00 ± 0.30 | -10.29 ± 0.07 | -10.69 ± 0.21 | -11.44 ± 0.43 |
| HJ16 | -10.76 ± 0.76 | -10.29 ± 0.07 | -10.13 ± 0.23 | -11.08 ± 0.21 |
| IOMA | -10.47 ± 0.27 | -9.12 ± 0.24 | -11.42 ± 0.34 | -10.52 ± 0.14 |
| J033 | -10.52 ± 0.13 | -8.63 ± 0.17 | -9.39 ± 0.19 | -10.29 ± 0.12 |
| J038 | -9.80 ± 0.21 | -9.04 ± 0.24 | -9.42 ± 0.19 | -9.87 ± 0.18 |
| N49P6 | -11.83 ± 0.29 | -10.44 ± 0.42 | -11.44 ± 0.30 | -10.83 ± 0.34 |
| N49P7 | -12.96 ± 0.42 | -12.35 ± 0.55 | -11.10 ± 0.38 | -11.73 ± 0.36 |
| N6-FR3-03 | -10.83 ± 0.27 | -9.65 ± 0.08 | -10.52 ± 0.11 | -10.37 ± 0.21 |
| N60P23 | -11.60 ± 0.31 | -10.99 ± 0.37 | -10.13 ± 0.17 | -11.35 ± 0.32 |
| NIH45-46 | -10.89 ± 0.36 | -10.20 ± 0.23 | -10.07 ± 0.17 | -11.05 ± 0.17 |
| PG9 | -9.97 ± 0.19 | -10.29 ± 0.07 | -9.49 ± 0.16 | -9.06 ± 0.12 |
| PGT121 | -10.40 ± 0.27 | -8.61 ± 0.28 | -10.81 ± 0.14 | -10.51 ± 0.15 |
| PGT122 | -10.15 ± 0.20 | -8.66 ± 0.20 | -9.56 ± 0.30 | -9.68 ± 0.34 |
| PGT128 | -9.04 ± 0.22 | -9.35 ± 0.20 | -9.54 ± 0.16 | -9.13 ± 0.18 |
| PGT145 | -10.07 ± 0.15 | -8.66 ± 0.17 | -9.26 ± 0.21 | -8.66 ± 0.13 |
| VH1-46 | -10.92 ± 0.21 | -10.29 ± 0.07 | -10.04 ± 0.27 | -10.39 ± 0.32 |
| VRC-CH31 | -11.48 ± 0.34 | -10.94 ± 0.37 | -11.31 ± 0.53 | -11.13 ± 0.23 |
| VRC-PG04 | -11.90 ± 0.18 | -11.88 ± 0.35 | -10.91 ± 0.29 | -12.12 ± 0.47 |
| VRC-PG20 | -10.96 ± 0.41 | -11.93 ± 0.31 | -11.51 ± 0.28 | -11.85 ± 0.40 |
| VRC01 | -12.58 ± 0.38 | -10.31 ± 0.06 | -11.63 ± 0.19 | -12.05 ± 0.31 |
| VRC01-523-LS-FR3-03 | -9.95 ± 0.36 | -9.73 ± 0.18 | -9.58 ± 0.27 | -10.51 ± 0.37 |
| VRC01-FR3-03 | -12.07 ± 0.25 | -10.20 ± 0.44 | -11.63 ± 0.42 | -11.26 ± 0.51 |
| VRC02 | -10.50 ± 0.15 | -11.21 ± 0.33 | -11.41 ± 0.23 | -11.60 ± 0.44 |
| VRC03 | -11.82 ± 0.16 | -10.29 ± 0.07 | -11.33 ± 0.44 | -11.80 ± 0.24 |
| VRC06b | -11.67 ± 0.54 | -11.24 ± 0.44 | -10.73 ± 0.30 | -11.05 ± 0.20 |
| VRC07 | -12.73 ± 0.33 | -12.46 ± 0.34 | -11.53 ± 0.34 | -10.52 ± 0.17 |
| VRC26.25 | -10.07 ± 0.24 | -8.94 ± 0.39 | -9.82 ± 0.16 | -9.45 ± 0.17 |

Table S4: Predicted dissociation constants ($K_d$, nM) of bNAbs across HIV-1 Env clades. Values are reported as mean ± STD. The expression $\Delta G = RT\ln K_d$ was used to get the dissociation constant of each bNAb, where R = 0.001987 kcal $mol^{-1}$ $K^{-1}$ and T = 300 K.

| bNAb | Clade A | Clade B | Clade C | Clade G |
|---|---|---|---|---|
| 10-1074 | 11.39 ± 3.60 | 41.30 ± 20.24 | 268.44 ± 79.65 | 30.64 ± 19.02 |
| 12A21 | 3.42 ± 3.66 | 3.24 ± 1.91 | 8.24 ± 5.23 | 14.65 ± 6.50 |
| 179NC75 | 196.27 ± 61.64 | 248.54 ± 139.67 | 43.28 ± 7.19 | 413.49 ± 134.43 |
| 1B2530 | 12.39 ± 5.38 | 17.79 ± 11.09 | 18.22 ± 5.46 | 27.35 ± 8.31 |
| 2411a | 4.46 ± 2.43 | 0.64 ± 0.37 | 5.55 ± 4.77 | 5.61 ± 2.92 |
| 2G12 | 167.17 ± 56.95 | 25.49 ± 3.78 | 111.16 ± 46.38 | 417.70 ± 123.16 |
| 35O22 | 6.25 ± 4.45 | 32.63 ± 3.73 | 7.65 ± 3.01 | 39.81 ± 6.74 |
| 3BNC117 | 0.22 ± 0.05 | 0.81 ± 0.58 | 1.24 ± 0.68 | 0.70 ± 0.16 |
| 3BNC55 | 3.24 ± 1.91 | 1.44 ± 0.36 | 1.30 ± 0.50 | 2.69 ± 1.55 |
| 438-B11 | 112.38 ± 53.40 | 570.61 ± 165.38 | 157.47 ± 50.22 | 37.18 ± 25.16 |
| 8ANC131 | 21.44 ± 7.44 | 3.65 ± 3.43 | 6.66 ± 2.76 | 4.23 ± 2.73 |
| b12 | 0.75 ± 0.43 | 1.53 ± 2.26 | 1.82 ± 1.88 | 504.47 ± 0.00 |
| b13 | 0.06 ± 0.01 | 32.22 ± 3.71 | 18.41 ± 1.73 | 0.07 ± 0.01 |
| BG24 | 8.84 ± 4.33 | 8.28 ± 10.12 | 6.69 ± 4.95 | 16.92 ± 6.43 |
| CAP257-RH1 | 56.15 ± 28.81 | 102.99 ± 45.58 | 43.35 ± 22.36 | 133.18 ± 41.01 |
| CH103 | 13.48 ± 7.69 | 214.67 ± 163.97 | 109.77 ± 26.68 | 3.26 ± 1.19 |
| CH235.12 | 28.04 ± 6.09 | 32.22 ± 3.71 | 40.83 ± 17.49 | 28.08 ± 9.95 |
| CH235.9 | 59.36 ± 159.88 | 32.25 ± 3.69 | 18.41 ± 1.73 | 504.47 ± 0.00 |
| DH270.5 | 238.51 ± 77.51 | 438.75 ± 142.10 | 29.20 ± 5.60 | 366.12 ± 62.11 |
| DH270.6 | 62.57 ± 31.67 | 74.15 ± 32.83 | 25.62 ± 6.76 | 504.47 ± 0.00 |
| DRVIA7 | 10.64 ± 4.36 | 31.98 ± 3.68 | 17.23 ± 6.62 | 5.78 ± 3.75 |
| HJ16 | 69.81 ± 198.94 | 32.22 ± 3.71 | 44.81 ± 18.60 | 8.93 ± 2.91 |
| IOMA | 25.86 ± 11.76 | 243.69 ± 93.69 | 5.66 ± 3.71 | 22.13 ± 5.32 |
| J033 | 22.24 ± 4.79 | 534.50 ± 166.31 | 149.63 ± 41.56 | 32.59 ± 6.19 |
| J038 | 76.85 ± 25.10 | 275.93 ± 88.21 | 142.56 ± 45.31 | 67.27 ± 27.00 |
| N49P6 | 2.64 ± 1.08 | 29.87 ± 16.88 | 5.14 ± 2.20 | 14.55 ± 6.18 |
| N49P7 | 0.45 ± 0.33 | 1.57 ± 1.90 | 9.52 ± 5.70 | 3.40 ± 2.24 |
| N6-FR3-03 | 13.97 ± 6.04 | 94.66 ± 12.77 | 21.98 ± 4.24 | 29.64 ± 13.01 |
| N60P23 | 4.00 ± 2.02 | 11.60 ± 6.85 | 42.94 ± 11.95 | 6.31 ± 4.52 |
| NIH45-46 | 13.65 ± 8.32 | 39.17 ± 11.69 | 47.98 ± 12.75 | 9.28 ± 2.61 |
| PG9 | 57.16 ± 19.34 | 31.98 ± 3.68 | 126.20 ± 35.93 | 256.46 ± 54.17 |
| PGT121 | 29.10 ± 13.81 | 597.80 ± 338.27 | 13.73 ± 3.22 | 22.59 ± 5.16 |
| PGT122 | 42.10 ± 13.16 | 517.66 ± 191.06 | 121.48 ± 69.27 | 102.20 ± 61.88 |
| PGT128 | 280.72 ± 145.96 | 160.80 ± 51.11 | 117.07 ± 35.25 | 232.29 ± 76.65 |
| PGT145 | 47.71 ± 11.86 | 511.15 ± 161.24 | 189.02 ± 58.55 | 501.65 ± 100.81 |
| VH1-46 | 11.82 ± 4.64 | 32.22 ± 3.71 | 53.31 ± 24.61 | 30.37 ± 15.70 |
| VRC-CH31 | 4.96 ± 2.63 | 12.89 ± 9.04 | 7.90 ± 5.73 | 8.42 ± 3.73 |
| VRC-PG04 | 2.21 ± 0.66 | 2.56 ± 1.52 | 12.49 ± 6.11 | 2.02 ± 1.91 |
| VRC-PG20 | 12.56 ± 8.26 | 2.33 ± 1.37 | 4.59 ± 2.58 | 2.87 ± 1.98 |
| VRC01 | 0.84 ± 0.64 | 31.16 ± 3.26 | 3.52 ± 1.22 | 1.86 ± 0.94 |
| VRC01-523-LS-FR3-03 | 73.00 ± 79.94 | 85.34 ± 27.71 | 117.05 ± 63.25 | 26.34 ± 17.24 |
| VRC01-FR3-03 | 1.75 ± 0.81 | 44.61 ± 23.30 | 4.21 ± 2.89 | 8.73 ± 7.88 |
| VRC02 | 22.87 ± 5.20 | 7.79 ± 4.28 | 5.19 ± 2.03 | 4.54 ± 3.38 |
| VRC03 | 2.54 ± 0.64 | 31.98 ± 3.68 | 6.96 ± 4.74 | 2.77 ± 1.48 |
| VRC06b | 4.55 ± 4.39 | 8.14 ± 5.39 | 16.70 ± 7.03 | 9.40 ± 2.83 |
| VRC07 | 0.62 ± 0.35 | 0.96 ± 0.54 | 4.67 ± 3.23 | 22.49 ± 6.78 |
| VRC26.25 | 49.94 ± 19.68 | 371.32 ± 233.74 | 72.68 ± 19.78 | 136.21 ± 36.82 |
